# Pyrophosphoproteomics: extensive protein pyrophosphorylation revealed in human cell lines

**DOI:** 10.1101/2022.11.11.516170

**Authors:** Jeremy A. M. Morgan, Arpita Singh, Leonie Kurz, Michal Nadler-Holly, Martin Penkert, Eberhard Krause, Fan Liu, Rashna Bhandari, Dorothea Fiedler

## Abstract

Reversible protein phosphorylation is a central signaling mechanism in eukaryotic cells. While the identification of canonical phosphorylation sites using mass-spectrometry (MS) based proteomics has become routine, annotation of non-canonical phosphorylation has remained a challenge. Here, we report a tailored pyrophosphoproteomics workflow to detect and reliably assign protein pyrophosphorylation in two human cell lines, providing the first direct evidence of endogenous protein pyrophosphorylation. Detection of protein pyrophosphorylation was reproducible, specific and consistent with previous biochemical evidence relating the installation of the modification to inositol pyrophosphates (PP-InsPs). We manually validated 148 pyrophosphosites across 71 human proteins, the most heavily pyrophosphorylated of which were the nucleolar proteins NOLC1 and TCOF1. A predictive workflow based on the MS data set was established to recognize putative pyrophosphorylation sequences, and UBF1, a nucleolar protein incompatible with the proteomics method, was biochemically shown to undergo pyrophosphorylation. When the biosynthesis of PP-InsPs was perturbed in a model cell line, proteins expressed in this background exhibited lower levels of pyrophosphorylation. Disruption of PP-InsP biosynthesis also significantly reduced rDNA transcription, potentially by lowering pyrophosphorylation on regulatory proteins NOLC1, TCOF1, and UBF1. Overall, protein pyrophosphorylation emerges as an archetype of non-canonical phosphorylation, and should be considered in future phosphoproteomic analyses.

## Introduction

The specific phosphorylation and dephosphorylation of proteins is a fundamental mechanism of intracellular signal transduction across the domains of life^1,2^. In humans, kinases and phosphatases dedicated to the writing and erasing of protein phosphorylation make up almost 2.5% of the genome^3–5^. After the discovery of serine (Fig. 1a), threonine, and tyrosine phosphorylation through biochemical approaches, mass spectrometry-based proteomics became the primary method to investigate the function and regulation of canonical Ser/Thr/Tyr phosphorylation^6^. As of 2022, more than 290,000 phosphorylation sites are reported in the PhosphositePlus database, identified almost entirely by phosphoproteomic approaches^7^.

**Figure 1.**
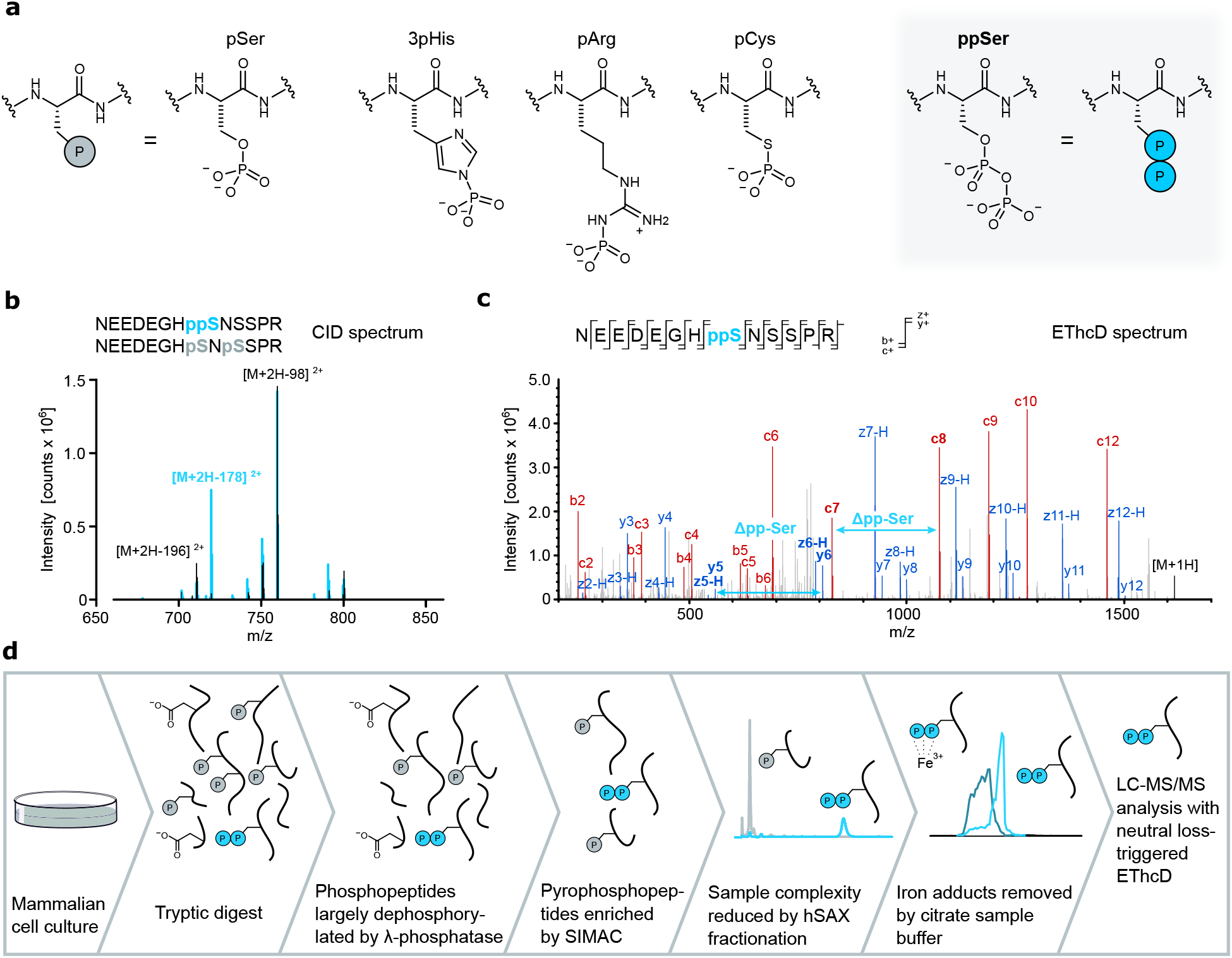
Enrichment and detection of pyrophosphorylated peptides using mass spectrometry. a) Examples of protein phosphorylation on different amino acid side chains. Canonical phosphorylation sites include phosphoserine (pSer), whereas non-canonical sites are exemplified by phosphohistidine (3-pHis), phosphoarginine (pArg), phosphocysteine (pCys), and pyrophosphoserine (ppSer). b) Fragmentation via collision induced dissociation (CID) of a pyrophosphopeptide results in a characteristic neutral loss (−178 Da), compared to the corresponding bisphosphopeptide. c) Fragmentation of a pyrophosphopeptide using electron-transfer dissociation combined with higher energy collision dissociation (EThcD) shows excellent sequence coverage, while leaving the modification intact. d) Development of a sample-preparation workflow, tailored to the enrichment and subsequent detection of tryptic pyrophosphopeptides.

In contrast, phosphorylation of non-canonical amino acid residues in humans (histidine, arginine, cysteine, aspartate, glutamate, lysine, Fig. 1a) has been established with varying success. Assignment of these sites was initially based on kinase assays utilizing radiolabeled ATP to mark phosphoryl acceptors for subsequent identification by autoradiography or radiochromatography, or later *via* immunodetection with antibodies^8^. These approaches are low-throughput and offer little site-specific information. Adapting traditional bottom-up phosphoproteomics workflows to allow for high throughput detection of these modifications has proved challenging, as the low pH conditions used to selectively enrich canonically phosphorylated peptides typically lead to the hydrolysis of the acid-labile phosphoramidate, thiophosphate, and acyl-phosphate moieties^9,10^. There is also an additional burden on the spectral interpretation, as convincing assignment of a novel non-canonical site requires the exclusion of a misassigned canonical site.

To address stability challenges, enrichment workflows with reduced or no dependence on low pH buffer systems have been developed, and include immunoprecipitation (pHis)^11^, online Fe^3+^ IMAC enrichment (pHis)^12^ and strong anion exchange (SAX) chromatography (pHis, pLys, pArg, pGlu, pCys, pAsp)^9^. To improve assignment accuracy, methods have incorporated characteristic neutral loss patterns^13^ and immonium ion formation^12^ into the detection and assignment process. In human backgrounds, identification of pHis is arguably the most advanced, with 14 sites biochemically validated^14^ and several hundred sites identified across different phosphoproteomics studies. However, there are still significant concerns regarding the validity of these high throughput site assignments^15^.

Protein pyrophosphorylation, the phosphorylation of a pSer residue to yield pyrophosphoserine (ppSer, Fig. 1a), is an additional non-canonical phosphorylation that is often overlooked^16–18^. This non-enzymatic posttranslational modification is mediated by high-energy inositol pyrophosphate messengers (PP-InsPs), which can transfer their β-phosphoryl group to protein substrates in the presence of Mg^2+^ ions^17^. Pyrophosphorylation was established using radiolabeled 5-diphosphoinositol pentakisphosphate (5PP-InsP_5_),the most abundant PP-InsP, demonstrating that a peptide or protein substrate can only accept the radiolabel at pre-phosphorylated residue^17^. ppSer exhibits differences in stability compared to pSer, notably a resistance to hydrolysis by common protein phosphatases and λ-phosphatase.

The *S. cerevisiae* proteins Nsr1, Srp40, and YGR130C were the first eukaryotic proteins shown to undergo pyrophosphorylation *in vitro*^16^. Subsequently, a handful of mammalian targets of *in vitro* protein pyrophosphorylation were identified and include nucleolar and coiled-body phosphoprotein 1 (NOLC1), Treacher Collins syndrome protein 1 (TCOF1), adaptor protein complex AP-3 subunit beta-1 (AP3B1), cytoplasmic dynein 1 intermediate chain 2 (DC1L2) and the oncoprotein MYC ^17,19–21^. Indirect evidence of endogenous pyrophosphorylation has relied on a “back-phosphorylation” assay, where potential targets are expressed and purified from a PP-InsP-rich cell line, and the subsequent *in vitro* phosphoryl transfer to this target is compared to the protein obtained from a cell line with low PP-InsP levels^22^. High PP-InsP levels will promote endogenous pyrophosphorylation, and therefore the subsequent *in vitro* phosphoryl transfer is decreased. While essential to discovery, these tools are low throughput and cannot provide direct information on the sites of modification.

Detection of endogenous pyrophosphorylation by mass spectrometry-based proteomics is an obvious solution to address these limitations. Compared to other modes of non-canonical phosphorylation, pyrophosphorylation is relatively acid-stable^16^, suggesting that traditional phosphoproteomic enrichment techniques should be compatible. However, the major technical challenge in enriching and detecting pyrophosphopeptides is their differentiation from peptides monophosphorylated at multiple positions. In particular, bisphosphorylated peptides (peptides containing two monophosphate groups) are isobaric, making the observation of the molecular ion uninformative. As with other non-canonical phosphorylation sites, incorrect assignment of canonical sites as non-canonical must be avoided^23^. In fact, unambiguous assignment requires an inversion of the traditional probability-based assignment; the peptide must be assumed to be bisphosphorylated, unless pyrophosphorylation can be asserted.

Here, we report the development of a dedicated pyrophosphoproteomics workflow for the detection and unambiguous assignment of endogenous pyrophosphorylation sites from mammalian cell lysates. Using neutral loss-triggered electron-transfer dissociation combined with higher energy collision dissociation (EThcD) LC MS/MS analysis, 108 and 78 sites were identified from HEK293T and HCT116 cell lines, respectively. Protein pyrophosphorylation sites predominantly occurred in acidic serine-rich stretches, and a majority of the identified pyrophosphoproteins localized to the nucleus and/or the nucleolus. In a functional readout for pyrophosphorylation of nucleolar proteins, we observed significantly impaired rDNA transcription in 5PP-InsP_5_-depleted cells. In sum, protein pyrophosphorylation can now be added unequivocally to the growing list of endogenous phosphorylation motifs in human cell lines.

## Results

### Establishment of a workflow for pyrophosphoproteomics

Using an array of synthetic peptide standards, we previously established that pyrophosphopeptides exhibit a characteristic neutral loss of –178 m/z, corresponding to the loss of pyrophosphoric acid (H_4_P_2_O_7_) (Fig. 1b)^24^. This neutral loss did not occur in Ser/Thr/Tyr monophosphorylated or bisphosphorylated peptides. Detection of this neutral loss could be used as a trigger during shotgun proteomic analysis to identify candidate pyrophosphopeptide ions, which then underwent EThcD fragmentation. The EThcD fragmentation spectra provided excellent sequence coverage while the modification stayed intact, and thus enabled identification of pyrophosphorylation sites (Fig. 1c, Fig. S1). With an optimized neutral loss filter, fine-tuned fragmentation parameters, and the exclusion of low-charge precursor ions, this triggered MS approach appeared suitable for the identification of pyrophosphorylation sites in cell lysates.

Therefore, a sample preparation workflow featuring enrichment of pyrophosphorylated tryptic peptides was developed (Fig. 1d). We tailored a standard phosphoproteomics workflow^25^ towards pyrophosphopeptide selection using a set of synthetic pyrophosphopeptides (and the corresponding phosphopeptides) of varying sequence characteristics for optimization (Fig. S2). Proteomic material was generated from HCT116 or HEK293T cells using standard protocols for cell culture and tryptic digestion. To reduce monophosphate competition during subsequent enrichment, the digested material was then treated with λ-phosphatase. This phosphatase hydrolyzed a large proportion of monophosphopeptides while leaving the pyrophosphoryl groups intact (Fig. S3)^17,26^. A SIMAC enrichment (sequential elution from immobilized metal ion affinity chromatography), featuring an additional low pH washing step designed to elute acidic peptides, was then performed^27^. Pyrophosphopeptides were largely retained during SIMAC (Fig. S4), while the overall material mass was reduced more than 40-fold. This retention is likely supported by the lower predicted pK_a_ value of the pyrophosphoryl moiety, in comparison to phosphoryl groups or acidic amino acid side chains.

Despite the phosphatase treatment and SIMAC enrichment, peptides with polyacidic amino acid stretches and multiple monophosphorylation sites were still abundant, so an offline fractionation step was implemented to further reduce sample complexity. A UPLC hydrophilic SAX (hSAX)^28^ column using a quaternary ammonium stationary phase on a hydrophilic polymeric support was selected, due to orthogonality with the low-pH reverse phase chromatography used in the LC-MS separation, and the ability to separate analytes of differing negative charge and polarity^6^. Separation was achieved using a buffer system with decreasing pH and acetonitrile gradients, and volatile reagents to maximize LC-MS compatibility^29,30^.

While establishing the workflow, we observed that pyrophosphorylated standard peptides were often detected in complex with Fe^3+^ ions during LC-MS analysis (Fig. S5). Crucially, these adducts formed in the liquid phase, as evidenced by distinct retention times and peak shapes. Such adduct formation reduced our ability to detect pyrophosphorylation sites, and thus sodium citrate (50 mM) was added into the sample resuspension buffer, which led to a substantial decrease of Fe^3+^ adduct formation (Fig. S5) ^31,32^. Overall, the sample preparation workflow (Fig. 1d) now seemed adequate for the enrichment of pyrophosphopeptides from complex samples.

### Reliable annotation of endogenous pyrophosphorylation sites

We next subjected the widely used mammalian cell line HEK293T to the pyrophosphoproteomics workflow. Using neutral-loss triggered EThcD mass spectrometry, many putative pyrophosphorylation sites were detected. To avoid incorrect assignment of multiply phosphorylated peptides (particularly bisphosphorylated peptides) as pyrophosphorylated, careful analysis of the data was required (Fig. 2a). Initial annotation was made on the basis of Sequest HT engine and ptmRS assignment as nodes in a Proteome Discoverer workflow. In the resulting data set, some spectra could be directly assigned as pyrophosphopeptides based on unambiguous fragmentation, but many spectra were annotated both as pyrophosphorylated and as bisphosphorylated peptides with comparable certainty. Increasing the threshold for the p-value during automated assignment did not alleviate this problem. It became apparent that co-elution (and co-fragmentation) of bisphosphorylated peptides with pyrophosphopeptides could produce ambiguous mixed spectra: if two bisphosphopeptides with overlapping phosphorylation pairs are co-fragmented, all fragments required to annotate a pyrophosphorylation site to that central residue are present (Fig. S7). This means the interpretation of each spectrum must exclude the possibility of such peptide mixtures before an assignment can be made. To accomplish this, a manual assessment process was developed.

**Figure 2.**
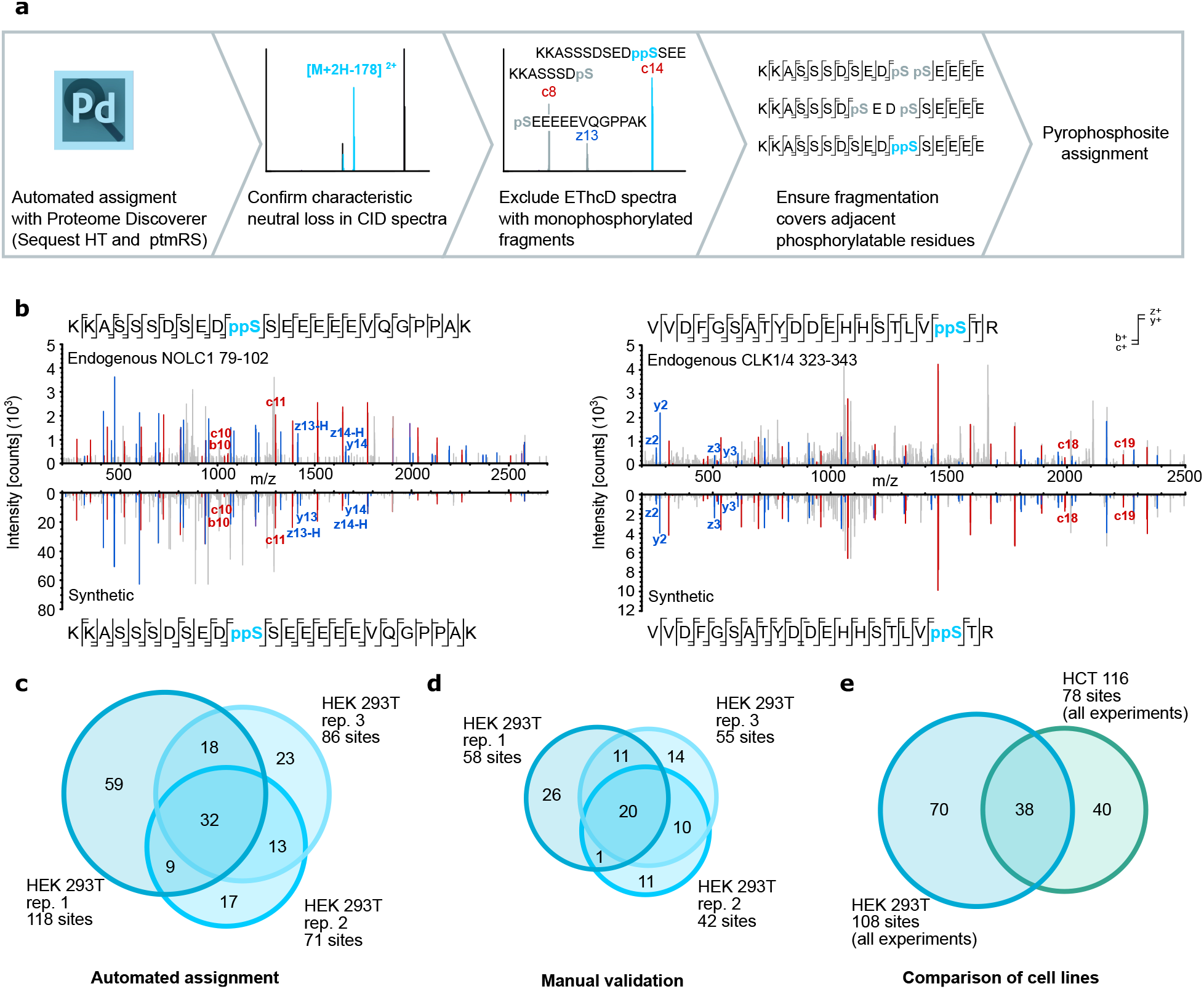
Assignment and validation of endogenous pyrophosphorylation sites. a) Workflow for the assignment of pyrophosphorylation sites following automated assignment with Proteome Discoverer. b) Comparison of EThcD spectra obtained from complex samples (top) with synthetically prepared pyrophosphopeptides (bottom). Fragment ions critical for the assignment of pyrophosphorylation sites are indicated in red (b- and c-ion series) or blue (y- and z-ion series). c) Venn diagram of the number of pyrophosphorylation sites detected by Proteome Discoverer in three biological replicates prepared from HEK293T cell lysates. d) Venn diagram of the number of pyrophosphorylation sites detected following manual assignment, using the three biological replicates from c). e) Overlap of pyrophosphorylation sites detected in HEK293T and HCT116 cells.

In the synthetic pyrophosphopeptide standards, the -178 m/z neutral loss peak was consistently among the three most intense signals in the CID spectra. The neutral loss was therefore selected as the criterion for identifying genuine pyrophosphorylated peptide ions, and candidate ions exhibiting -178 m/z neutral loss signals outside of the three highest were discarded. The EThcD fragmentation spectra of candidate ions were then manually examined using Molecular Weight Calculator (https://github.com/PNNL-Comp-Mass-Spec/Molecular-Weight-Calculator-VB6)^33^ for evidence of peptide fragments containing a single phosphorylated residue - this is only possible if the peptide is bisphosphorylated. Again, candidate ions exhibiting such fragments were discarded (Fig. 2a). Finally, the extent of fragmentation across the putative pyrophosphorylation site was assessed. Missing fragments, particularly those encompassing a canonically phosphorylatable residue such as serine or threonine can lead to the misassignment of a bisphosphorylated peptide as pyrophosphorylated, and as such, spectra with key fragments missing were discarded. The remaining spectra correspond to genuine pyrophosphorylation sites (Fig. 2a).

To validate the pyrophosphosite assignment, we synthesized pyrophosphopeptides based on two detected sequences, NOLC1 79-102 and CLK1/4 323-343 (Fig. 2b). Both peptide sequences contain phosphorylatable residues immediately adjacent to the putative pyrophosphorylation sites. To prove that pyrophosphorylation is present, sequential ions consistent with the unphosphorylated peptide fragment immediately preceding the putative site (e.g. ppNOLC1 79-88) and the sequential pyrophosphorylated peptide fragment containing the putative site (e.g. ppNOLC1 79-89) must be detected simultaneously (in the absence of a singly phosphorylated peptide fragment indicative of bisphosphorylation). For the NOLC1 sequence, the c10 and c11 ion couplet and the z13-H and z14-H ion couplet in the c/z ions series were observed in both the fragmentation spectra of the synthetic peptide and of the endogenous peptide, confirming the presence of the putative pyrophosphorylation site. Similarly, the CLK1/4 sequence exhibited the z2/z3 and c18/c19 ion couplets in the c/z ions series, and the y2/y3 ion couplet in the b/y ion series, in both synthetic and endogenous peptide fragmentation spectra, consistent with pyrophosphorylation at Ser341. No fragments indicative of monophosphorylation were detected in any spectra and crucially, the diagnostic ion couplets were absent in the fragmentation spectra of the corresponding bisphosphopeptide (Fig. S6). Together, these data validated our assignment approach for the reliable and correct identification of endogenous pyrophosphorylation sites.

After applying the pyrophosphoproteomics workflow and assignment strategy to HEK293T cell lysates, three biological replicates were submitted to the analysis. Each sample was injected twice, and the identified sites combined. A total of 171 unique pyrophosphorylation sites were detected by automated assignment, 93 of which were manually validated (an average of 51 per replicate, Fig. 2c,d). 40 pyrophosphoproteins were identified from manually validated sites. While a core of 20 manually validated sites were seen in all replicates, a significant portion of the sites (51) were exclusively observed in a single replicate. This is likely due to variation in the sample background of each replicate, differentially co-eluting with, and masking these low abundant species in replicates where they are not detected. Three proteins exhibiting multiple pyrophosphorylation sites were heavily overrepresented in the triplicate; NOLC1, TCOF1 and SRRM1 (serine/arginine repetitive matrix protein 1) sites were found in all replicates and represented 43 of the 93 total sites assigned.

During method development, lysates from another human cell line, HCT116 colon cancer cells, were also frequently subjected to pyrophosphoproteomic analysis and a total of 78 sites on 33 proteins were identified (Table S1). Many of these pyrophosphorylation sites overlap with sites from HEK293T lysates (Fig. 2e), further corroborating the reliability of the assignment strategy.

### Protein pyrophosphorylation is commonly primed by CK2 and detected in nuclear and nucleolar proteins

Two proteins found to be heavily pyrophosphorylated were the nucleolar proteins NOLC1 and TCOF1, containing 34 and 18 different pyrophosphorylation sites, respectively. These observations are consistent with previous biochemical studies, in which both NOLC1 (also called Nopp140) and TCOF1 were able to undergo *in vitro* radiolabeling by [β^32^P]5PP-InsP_5_^16,17^. Both NOLC1 and TCOF1 are densely phosphorylated proteins, which presumably facilitates their pyrophosphorylation. Interestingly, the pyrophosphorylation sites on NOLC1 and TCOF1 exclusively localize to acidic regions, whereas the phosphorylation sites are reported to be more evenly distributed across acidic and basic regions (Fig. 3a). The localization of pyrophosphorylation sites to acidic serine stretches has been observed in previous studies on 5PP-InsP_5_-mediated pyrophosphorylation^17,19–21,34,35^. In all cases, the priming *in vitro* phosphorylation was catalyzed by acidophilic Ser/Thr kinases, especially CK2. A global analysis of all endogenous pyrophosphorylation sites detected by mass spectrometry in HEK293T and HCT116 lysates confirmed the central role for acidophilic Ser/Thr kinases - alignment of all sequences revealed a clear CK2 consensus sequence (Fig. 3b)^36^. However, some pyrophosphorylation sequences did not match the CK2 consensus sequence. When we removed all sites containing a Glu/Asp/Ser/Thr residue in position +3 from the alignment, a proline-directed kinase consensus sequence emerged^37^ (Fig 3b), suggesting that this family of Ser/Thr kinases may also pre-phosphorylate residues prior to their pyrophosphorylation.

**Figure 3.**
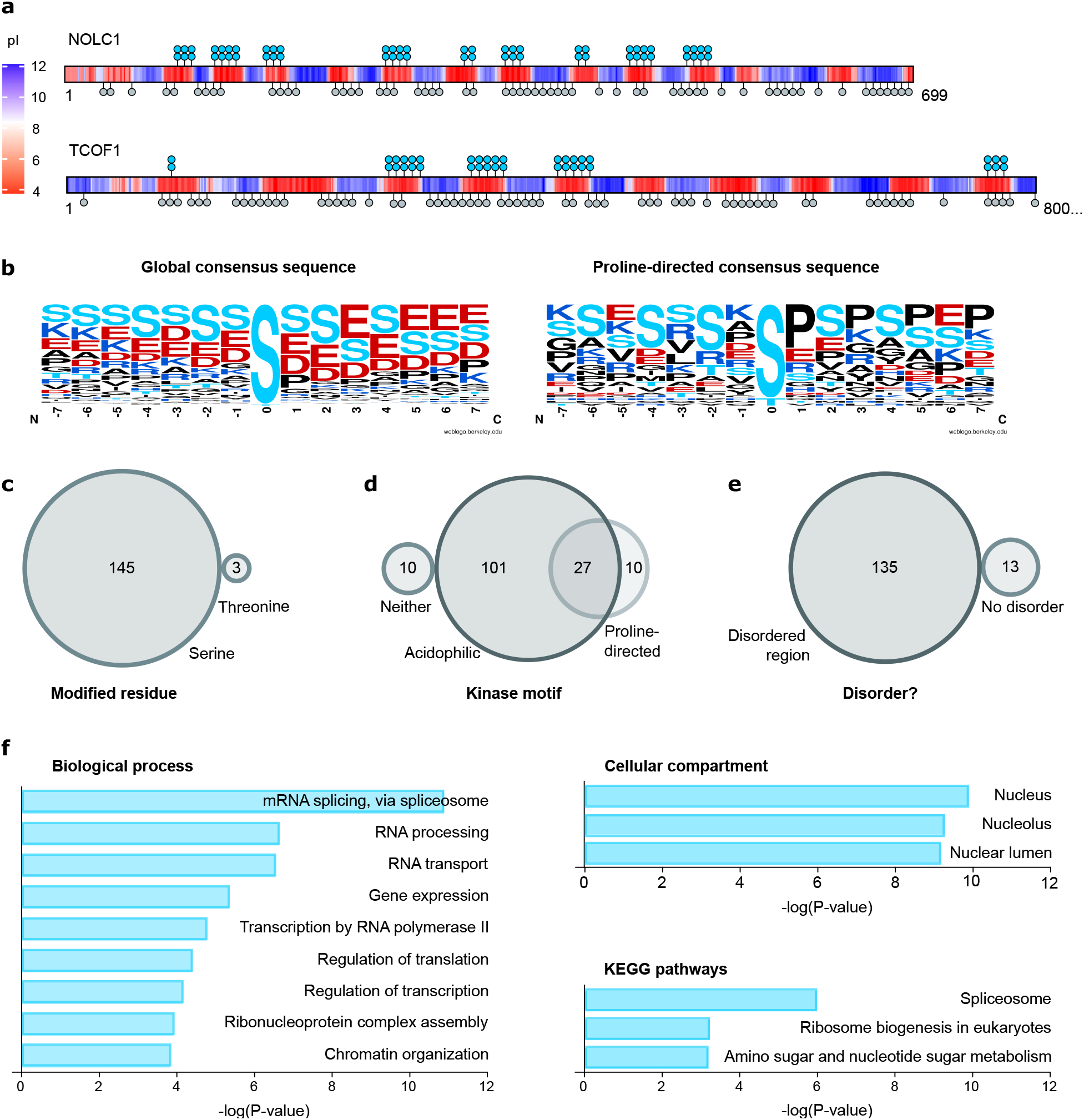
Properties of pyrophosphorylation sites. a) Illustration of the localization of pyrophosphorylation sites to acidic regions in NOLC1 and TCOF1. The local pI was calculated using a custom R-script. Pyrophosphoserine residues are shown in turquoise, and previously reported phosphorylation sites are indicated in grey (Phosphosite Plus® ^7^). b) Consensus sequence of all pyrophos-phorylation sites detected (left), and consensus sequence following removal of CK2 consensus sites (right). Sequence logos were generated using WebLogo^41^ c) Side-chain of modification for all pyrophosphorylation sites. d) Kinase motifs surrounding the pyrophosphorylation sites. e) Analysis of order/disorder around pyrophosphorylation sites. f) Gene ontology analysis of pyrophosphoproteins using Enrichr (Table S3, https://maayanlab.cloud/Enrichr/)^42–44^.

Of the 148 pyrophosphorylated sites identified in our study, only three occur on threonine residues, and the remainder are on serine (Fig. 3c). We analyzed the pyrophosphosites using the Scansite 4.0 tool, to predict motifs that are likely to undergo phosphorylation by specific protein kinases (https://scansite4.mit.edu/#scanProtein)^38^. Consistent with the consensus sequences above, a majority of the sites were predicted to be phosphorylated by acidophilic Ser/Thr kinases, a smaller number were potential substrates for proline-directed Ser/Thr kinases, and a few sites may be substrates for both families of kinases (Fig. 3d). Approximately 7% of the mapped pyrophosphorylated residues were not predicted as substrates for either acidophilic or proline-directed Ser/Thr kinases, suggesting that other families of protein kinases may also prime residues for pyrophosphorylation. An additional feature common to pyrophosphorylation sites is that they lie within intrinsically disordered regions (IDRs)^18^. A disorder prediction analysis showed that 91% of the pyrophosphosites identified in our study lie within a continuous stretch of 20 or more residues that have a disorder score ≥ 0.5, which is the cutoff for predicted disorder at that residue (Fig. 3e). This is in line with properties of phosphorylation sites in general, as IDRs are overrepresented in eukaryotic phosphoproteomic datasets^39,40^.

Gene ontology analysis of pyrophosphorylation sites suggests a function for this modification in nuclear and nucleolar processes, as these two compartments were significantly overrepresented among pyrophosphorylated proteins (Table S3, Fig. 3f). This localization preference is mirrored by the biological processes, in which RNA processing, specifically RNA splicing, emerged as a process putatively regulated by protein pyrophosphorylation of SRRM1, SRRM2 (Serine/arginine repetitive matrix protein 1,2), SRSF2, SRSF5, SRSF6, SRSF9 (Serine/arginine-rich splicing factor 2,5,6,9), SF3B2 (Splicing factor 3B subunit 3), WBP11 (WW domain-binding protein 11), and TRA2B (Transformer-2 protein homolog beta), among others. Proteins NOLC1, NOP58 (nucleolar protein 58), NPM1 (nucleophosmin), DKC1 (H/ACA ribonucleoprotein complex subunit DKC1), and MPHOSPH10 (U3 small nucleolar ribonucleoprotein protein MPP10) are involved in ribosome biogenesis, another process overrepresented among pyrophosphoproteins (Fig. 3f). A functional connection between PP-InsPs and ribosome biogenesis had previously been made in *S. cerevisiae* using genetics, and a mechanistic hypothesis for this regulation involved pyrophosphorylation of the RNA polymerase I subunits A190, A43 and A34.5 ^35^.

### The nucleolar fibrillar center is enriched in pyrophosphorylated proteins

The nucleolus, which emerged as a major site for localization of pyrophosphorylated proteins, is a membrane-less organelle organized into three liquid-liquid phase-separated sub-compartments – the fibrillar center (FC), dense fibrillar component (DFC), and granular component (GC) (Fig. 4a)^45^. rDNA repeats present in the FC are transcribed by RNA polymerase I at the FC/DFC boundary; the resulting pre-rRNAs are processed in the DFC and assembled into ribosomes in the GC^45^. Human IP_6_ kinase isoforms IP6K1 and IP6K2 are reported to be localized to the nucleolar FC region (Human Protein Atlas, https://www.proteinatlas.org/)^46^. We used immunofluorescence to confirm the co-localization of IP6K1 with the FC marker protein UBF1 (Upstream binding factor 1) in HEK293T cells (Fig. 4b). The same pattern of localization was observed in the osteosarcoma cell line U-2 OS (Fig. 4b). Super-resolution microscopy revealed that IP6K1 is confined to the FC and does not co-localize with FBL (fibrillarin) in the DFC (Fig. 4c,d). Local synthesis of 5PP-InsP_5_ by IP6Ks in the FC would facilitate pyrophosphorylation of FC resident proteins. Indeed, the two most highly pyrophosphorylated proteins identified in our study, NOLC1 and TCOF1, are localized to the FC. Four additional pyrophosphoproteins - NOP58, DKC1, SMARCA4 (SWI/SNF related, matrix associated, actin dependent regulator of chromatin, subfamily a, member 4), and EEF1D (eukaryotic translation elongation factor 1 delta) are also annotated to the FC. Other proteins enriched in this nucleolar compartment thus appear likely targets of pyrophosphorylation. As most pyrophosphorylated sequences fall within IDRs, and can potentially be pre-phosphorylated by acidophilic and/or Pro-directed Ser/Thr kinases (Fig 3d and e), these criteria were used to uncover additional FC localized substrates for pyrophosphorylation. We used IuPred2A (https://iupred2a.elte.hu/)^47^ to predict regions of disorder, and Scansite 4.0 (https://scansite4.mit.edu/#scanProtein)^38^ to predict pre-phosphorylation sites on 144 human proteins annotated to the cellular component gene ontology term ‘fibrillar center’ (http://amigo.geneontology.org/amigo/landing)^48^ (Fig. 4e). Of these, 100 FC annotated proteins were predicted to possess potential sites for pyrophosphorylation, including the six FC proteins identified by mass spectrometry (Table S4), suggesting that our pyrophosphoproteomic workflow may not discover all pyrophosphoproteins.

**Figure 4.**
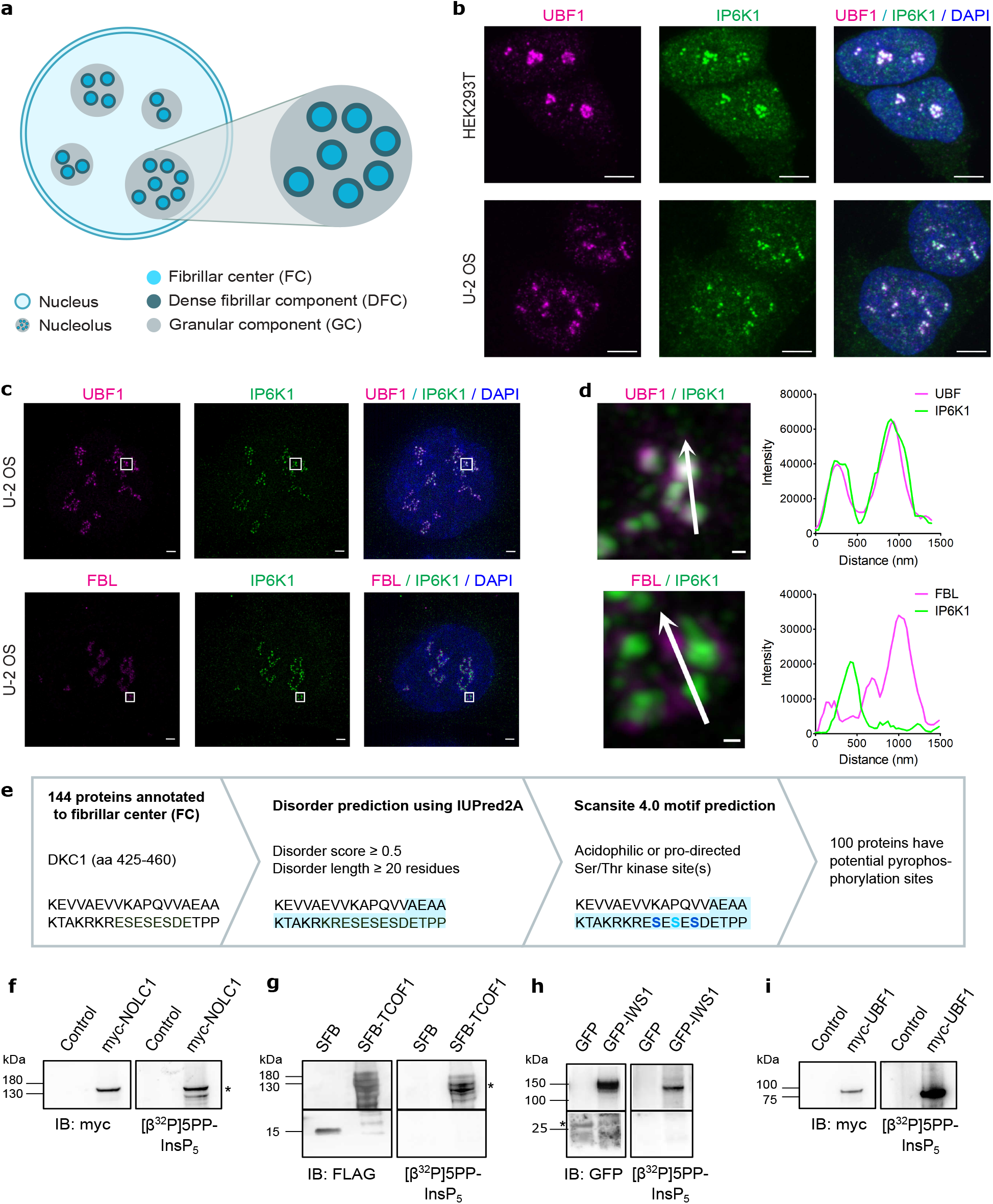
Nucleolar FC proteins undergo 5PP-InsP_5_-mediated pyrophosphorylation. a) Architecture of the mammalian nucleolus and its sub-compartments. b) Confocal micrographs of HEK293T and U-2 OS cells showing co-localization of IP6K1 (green) with nucleolar FC marker UBF1 (magenta). Nuclei were stained with DAPI (blue); scale bars are 5 μm. c) Super-resolution (SIM) images of IP6K1 (green) colocalized with UBF1 or the nucleolar DFC marker FBL (magenta) in U-2 OS cells. Nuclei were stained with DAPI (blue); scale bars are 2 μm. d) Magnification of the boxed region in (c); scale bars are 0.2 μm. Traces show fluorescence intensity profiles for IP6K1 (green) and UBF1 or FBL (magenta), measured along the indicated arrow. e) Workflow to predict potential pyrophosphorylation sites in 144 human proteins annotated to the nucleolar FC, using human DKC1 (residues 425-460) as an example. The light blue area indicates a predicted IDR; Ser residues in blue and turquoise are predicted and identified pyrophosphosites, respectively. f-i) *In vitro* pyrophosphorylation of human proteins by [β^32^P]5PP-InsP_5_. NOLC1, TCOF1, IWS1 and UBF1 with their indicated N-terminal tags were expressed in HEK293T cells, isolated, and prephosphorylated by CK2 prior to incubation with [β^32^P]5PP-InsP_5_. Images show autoradiography to detect pyrophosphorylation (right) and immunoblotting with respective tag-specific antibodies (left). Negative controls were cells transfected with plasmids pCMV-myc (f), pCDNA-SFB (g) or EGFP-C1 (h), or untransfected cells (i). The asterisks in f, g and h indicate specific bands.

Therefore, we used radiolabeled [β^32^P]5PP-InsP_5_ as an alternative method to test for pyrophosphorylation of candidate proteins. We confirmed that pyrophosphoproteins identified by mass spectrometry - NOLC1, TCOF1, and the nuclear protein IWS1 – can undergo *in vitro* pyrophosphorylation with [β^32^P]5PP-InsP_5_ following pre-phosphorylation by CK2 (Fig. 4f-h). UBF1, which co-localizes with IP6K1 in the FC (Fig. 4b-d), is predicted to possess sites for pyrophosphorylation (Table S4). CK2 pre-phosphorylation and exposure to [β^32^P]5PP-InsP_5_ brought about robust *in vitro* pyrophosphorylation of overexpressed UBF1 isolated from HEK293T cells (Fig. 4i). A comparison of the predicted pyrophosphorylation sites on UBF1 with the mapped pyrophosphorylated sites on NOLC1 (Fig. S8) revealed a difference between the properties of their neighboring sequences. Unlike the pyrophosphoserine residues on NOLC1, the predicted UBF1 pyrophosphosites lie either within very short or very long tryptic fragments, both of which would evade detection by our current pyrophosphoproteomic workflow.

### PP-InsPs are responsible for intracellular pyrophosphorylation and support rDNA transcription

Given the complementarity of the radiolabeling and mass spectrometry methods, we used both approaches to affirm that 5PP-InsP_5_ drives intracellular protein pyrophosphorylation. We relied on *IP6K1*^*-/-*^ HEK293T cells expressing either active or kinase-dead IP6K1 along with TCOF1, NOLC1 or IWS1 (Fig. 5a, Fig. S9)^49^. These substrate proteins were purified, treated with λ-phosphatase, and resolved by SDS-PAGE; the excised gel pieces were digested with trypsin and measured by neutral loss-triggered EThcD MS as described above. Equal loading between conditions was confirmed by comparing the sum of CID precursor intensities (Fig. S10). There were many neutral-loss triggers generated by the characteristic loss of the pyrophosphoryl moiety in the samples obtained from cells expressing active IP6K1, several of which could be confirmed to correspond to pyrophosphorylated peptides for all three substrates (Fig. 5a and Table S5). By contrast, we could not identify a single peptide sequence triggered by the characteristic loss of the pyrophosphoryl moiety from the cells expressing kinase-dead IP6K1. These results illustrate the strong dependence of endogenous pyrophosphorylation on cellular PP-InsP levels.

**Figure 5.**
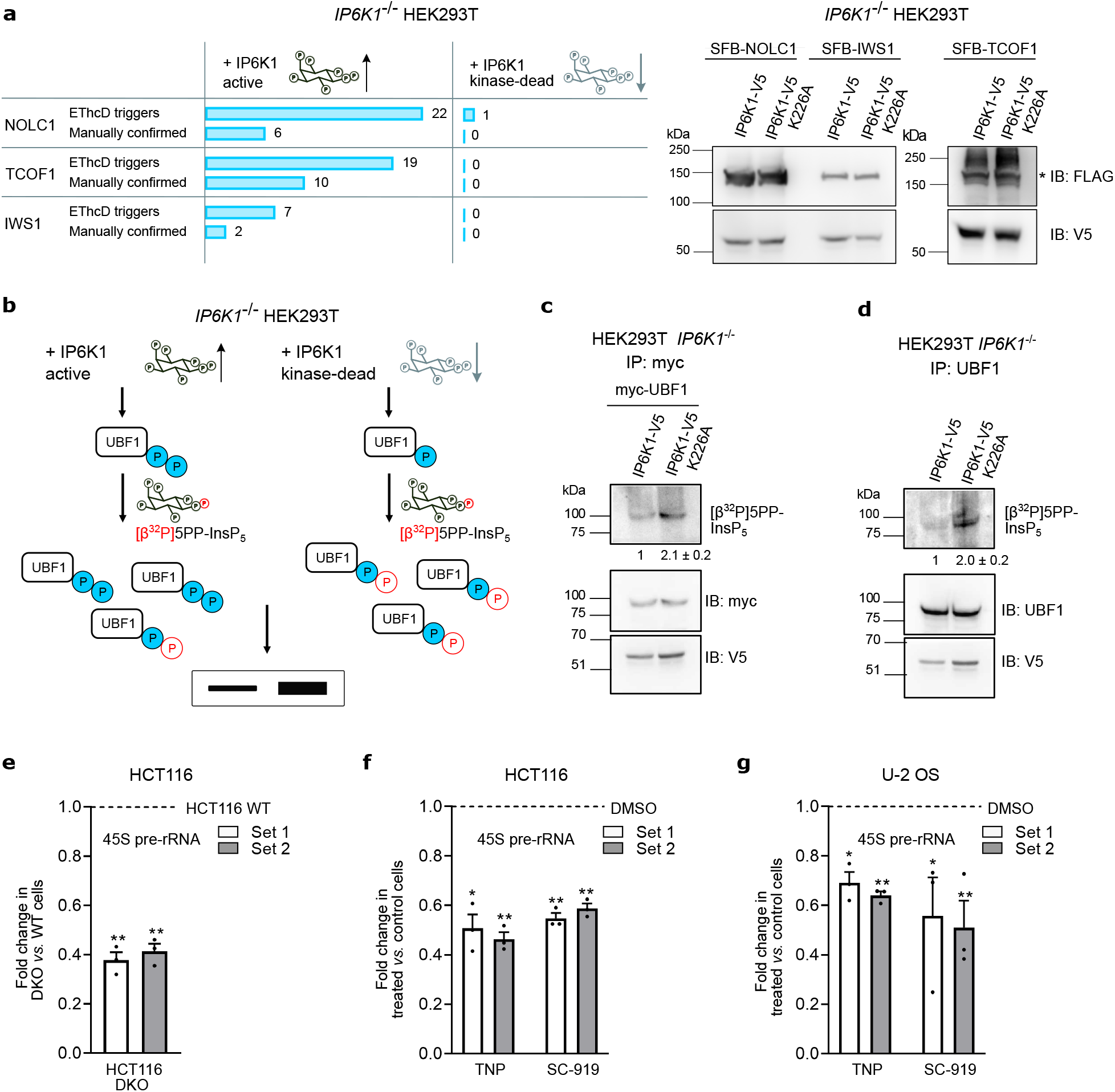
5PP-InsP_5_ drives cellular pyrophosphorylation and promotes rRNA synthesis. a) Comparative MS analysis of pyrophosphosites detected in SFB-tagged NOLC1, TCOF1 or IWS1 co-expressed with V5-tagged active or kinase-dead IP6K1 (left). Number of neutral-loss triggered EThcD spectra vs pyrophosphopeptides confirmed by manual analysis criteria (Fig.2). Immunoblot shows that equal amounts of protein were subjected to MS (right). Asterisk indicates the specific band. (b) Back-pyrophosphorylation method to detect intracellular UBF1 pyrophosphorylation. c)-d) Overexpressed myc-tagged UBF1 (c) or endogenous UBF1 (d) were immunoprecipitated from *IP6K1*^-/-^ HEK293T cells expressing either active or kinase-dead V5-tagged IP6K1, and incubated with [β^32^P]5PP-InsP_5_. Images show autoradiography to detect pyrophosphorylation (top) and immunoblotting with the indicated antibodies (bottom). Numbers show mean fold change ± SEM in the extent of UBF1 back-pyrophosphorylation in cells expressing kinase-dead compared to active IP6K1 (N=3 (c) and 4 (d)). e-g) RT-qPCR analysis to measure 45S pre-rRNA transcript levels using two different primer sets. Values indicate the fold change in transcript levels in *IP6K1*^*-/-*^ *IP6K2*^*-/-*^ double knockout (DKO) cells compared to HCT116 WT cells (e), and TNP or SC-919 treated HCT116 (f) or U-2 OS (g) cells compared to cells treated with the vehicle control (DMSO). Data are mean ± SEM (N=3). *P*-values were determined using a one-sample t-test. \*\**P* ≤ 0.01 and \**P* ≤ 0.05.

As pyrophosphosites on UBF1 evade detection by mass spectrometry, we resorted to the ‘back-pyrophosphorylation’ method to examine intracellular UBF1 pyrophosphorylation^22^ (Fig. 5b). Overexpressed or endogenous UBF1 was isolated from *IP6K1*^*-/-*^ HEK293T cells expressing either active or kinase-dead IP6K1, and incubated with radiolabeled [β^32^P]5PP-InsP_5_. UBF1 isolated from cells expressing kinase-dead IP6K1 showed a two-fold higher pyrophosphorylation signal on the autoradiogram compared with UBF1 from cells with active IP6K1, reflecting higher levels of intracellular pyrophosphorylation on the latter form of UBF1 (Fig. 5c,d). The increase in cellular pyrophosphorylation on TCOF1, NOLC1, IWS1 and UBF1 in the presence of active IP6K1 is consistent with intracellular pyrophosphorylation mediated by 5PP-InsP_5_.

TCOF1, NOLC1 and UBF1 are known regulators of RNA polymerase I-mediated rDNA transcription^43–46^. Reduced intracellular 5PP-InsP_5_, which in turn will lower pyrophosphorylation levels of these proteins, is therefore likely to have an impact on rRNA synthesis. To examine this possibility, we utilized two different cellular models known to be depleted for 5PP-InsP_5_ - (a) HCT116 cells lacking IP6K1 and IP6K2^54^ and (b) cells treated with the InsP_6_ kinase inhibitors TNP or SC-919^55,56^. Quantitative RT-PCR analysis was used to assess the levels of the 45S pre-rRNA transcript, which is subsequently processed to yield mature rRNA for incorporation into ribosomes. We observed a more than two-fold decrease in the levels of 45S pre-rRNA in *IP6K1*^*-/-*^ *IP6K2*^*-/-*^ double knockout (DKO) cells compared to wild type HCT116 cells. Treatment with IP6K inhibitors recapitulated these findings, and similarly suppressed pre-rRNA synthesis in HCT116 and U-2 OS cells, compared to cells treated with the vehicle control. These observations point to a significant role for 5PP-InsP_5_ in the maintenance of rDNA transcription, likely *via* pyrophosphorylation of key nucleolar proteins.

## Discussion

Here, we have developed a tailored pyrophosphoproteomics workflow to detect and assign protein pyrophosphorylation in two human cell lines (148 manually validated sites across 71 proteins), providing the first direct evidence of endogenous protein pyrophosphorylation. Over time, efforts to characterize non-canonical phosphorylation with phosphoproteomic methods have advanced, as sample handling, enrichment and mass spectrometry techniques have been adapted to the properties of the modification. For example, pHis proteomics has evolved by leveraging antibody enrichment and later strong anion-exchange enrichment, culminating in the establishment of a dedicated pHis database, HisPhosSite, containing more than 270 human sites.^57^ Despite these advances, two major technical challenges have remained: Site localization and therefore distinction from canonical phosphorylation is difficult, and as site localization probabilities are tightened in assignment workflows, unambiguous assignments are dramatically reduced^9^. Lemeer and coworkers recently used the pHis immonium ion as additional evidence of correct pHis assignment, but in their human dataset, only 0.5% of putative pHis-containing peptides exhibited this ion, suggesting significant overassignment by current automated methods^15^. Moreover, pHis and N-phosphorylated peptides in general have been shown to undergo phosphoryl-transfer reactions in the gas-phase, changing the site of modification prior to characterization^58^.

In the case of pyrophosphorylation, we observed that the challenges of acid lability and potential gas phase transfer associated with N-phosphorylation were absent. With an appropriately designed workflow and enrichment method, generating sufficient sequence coverage across the putative site upon EThcD fragmentation, we could achieve unambiguous pyrophosphosite assignment. The proteomic datasets reproduced the trends observed in previous biochemical characterization; pyrophosphorylation occurs in acidic regions, often in stretches known to be multiply phosphorylated. This meant that mixtures of isobaric bisphosphopeptide isomers and pyrophosphopeptides are likely co-fragmented during EThcD, increasing the risk of false assignment. To exclude this possibility, both the CID and EThcD fragmentation patterns needed to be assessed, and spectra containing monophosphorylated fragments were excluded from the analysis. While this manual workflow is more laborious than automated methods, the assigned pyrophosphorylation sites are unambiguous, and can be directly used to inform further biological and biochemical investigation. On this basis, we conclude that pyrophosphorylation may be the most abundant non-canonical phosphorylation characterized in human cells to date.

To demonstrate the dependence of protein pyrophosphorylation on PP-InsP levels, we qualitatively assessed the difference in the number of characteristic neutral loss triggers during CID fragmentation of NOLC1, TCOF1 and IWS1 expressed in cells with high (IP6K1-kinase active) or low (IP6K1-kinase dead) PP-InsP levels. Substantially more triggers were observed for all proteins expressed in the IP6K1-kinase active background (48 *versus* 1 across all proteins, Fig. 5a), and not a single pyrophosphorylation site could be assigned in the IP6K1-kinase dead background. All three proteins were also able to accept the radiolabeled phosphoryl group from [β^32^P]5PP-InsP_5_, demonstrating that the pyrophosphorylation of these proteins is consistent with the non-enzymatic model of protein pyrophosphorylation.

In the future, a quantitative mass spectrometry approach (such as TMT labeling or SILAC) should be developed, so that levels of protein pyrophosphorylation in different cellular backgrounds, or under different conditions, can be compared. Other elements of the workflow would also benefit from further optimization, for example, to reduce the amount of material required per analysis, and to improve the level of overlap between biological replicates. Both issues likely stem from the limitations of the enrichment step efficiency, where non-pyrophosphorylated material remains after enrichment and fractionation and then stochastically masks pyrophosphopeptide ions during LC separation and CID fragmentation. Implementation of online-Fe-IMAC chromatography^59^ might allow for improved selectivity by fine-tuning pH and elution conditions, reducing material requirements and increasing reproducibility. Additionally, the lack of Lys/Arg residues for tryptic cleavage in acidic polyserine stretches may prevent detection of pyrophosphorylated peptides by our approach. In the future, this limitation could be addressed by integrating additional proteases into the digestion step, together with trypsin, such as GluC which cleaves at the C-terminus of aspartic acid residues^60^.

NOLC1 and TCOF1 were the two most heavily pyrophosphorylated proteins identified in this study. Along with UBF1, which we characterized as a novel *in vitro* pyrophposphorylation substrate, NOLC1 and TCOF1 are known to upregulate rRNA synthesis via RNA Pol I binding^50–53^. CK2-dependent phosphorylation at the C-terminal region of UBF1 promotes its interaction with SL1, to form a stable RNA Pol I preinitiation complex^53,61^. As most of the predicted UBF1 pyrophosphorylation sites lie within its C-terminus (Fig. S9), pyrophosphorylation on UBF1 subsequent to its pre-phosphorylation by CK2 may conceivably regulate rRNA synthesis. Consistent with this, depletion of 5PP-InsP_5_ leads to decreased rDNA transcription (Fig. 5e-g), likely via Pol I regulation. We have earlier shown that in the budding yeast *Saccharomyces cerevisiae*, the absence of 5PP-InsP_5_ leads to severely reduced rRNA synthesis, correlating with pyrophosphorylation of RNA Pol I subunits^35^. 5PP-InsP_5_ is thought to be a ‘metabolic messenger’ as its synthesis by IP6Ks is uniquely sensitive to ATP availability^62,63^. If the interaction between NOLC1/TCOF1/UBF1 and RNA Pol I were indeed dependent on 5PP-InsP_5_-mediated pyrophosphorylation, this would represent a straightforward cellular energy sensing mechanism to control rDNA transcription.

The localization of pyrophosphoproteins to the nucleolus was quite striking and raises the question whether pyrophosphorylation plays a general regulatory role in this biomolecular condensate. A recent study showed that phosphorylation of specific sites influenced partitioning of NPM1 (pSer125) and HNRNPA1 (Heterogeneous nuclear ribonucleoprotein A1, pSer6) to the nucleolus^64^. Interestingly, in our MS data both proteins were found to be pyrophosphorylated at these specific positions. It would therefore be desirable to develop tools that can accurately represent or mimic pyrophosphorylation at the protein level. Such tools would enable researchers to investigate the influence of pyrophosphorylation on protein partitioning into membraneless condensates.

Overall, protein pyrophosphorylation has emerged as an abundant, non-canonical phosphorylation. The ability to detect this modification within complex samples using MS, and to definitively assign the modification sites, now opens the door to investigate the regulatory role of protein pyrophosphorylation, both at the biochemical and cellular level. While the installation of pyrophosphorylation appears to be non-enzymatic in biochemical assays, it is still an open question whether the presence of cell lysates or other co-factors can accelerate protein pyrophosphorylation. How the features of the PP-InsP phosphoryl donor influence the degree and the specificity of pyrophosphorylation has also not been addressed to date. Following installation, it appears feasible that pyrophosphorylation could be detected by specific reader domains, and knowledge of the pyrophosphoproteome will facilitate the identification of such readers. And finally, the removal of pyrophosphorylation sites will need to be explored. Are there dedicated protein pyrophosphatases that convert pyrophosphoserine back to phosphoserine or serine? Elucidating these questions will provide fundamental insight into the regulation of protein pyrophosphorylation, and its interplay with signaling pathways controlled by canonical protein phosphorylation.

## Acknowledgements

*The authors thank the FMP peptide facility (Ines Kretzschmar) for assistance with peptide synthesis and Boris Bodganov for providing the R script used in Figure 3*.

*We also thank the CDFD microscopy facility for their support and Shubhra Ganguli, Jayashree Ladke, Akruti Shah, Ruth Manorama for HPLC analysis and generation of IP6K1 KO cell lines*.

Dedicated to Katherine M. Hannan, for demonstrating that the most important trait a researcher can have is kindness.

## Notes

### Competing Interest Statement

The authors have declared no competing interest.

