## Supplementary material for "Pyrophosphoproteomics: extensive protein pyrophosphorylation revealed in human cell lines": Methods and supporting figures

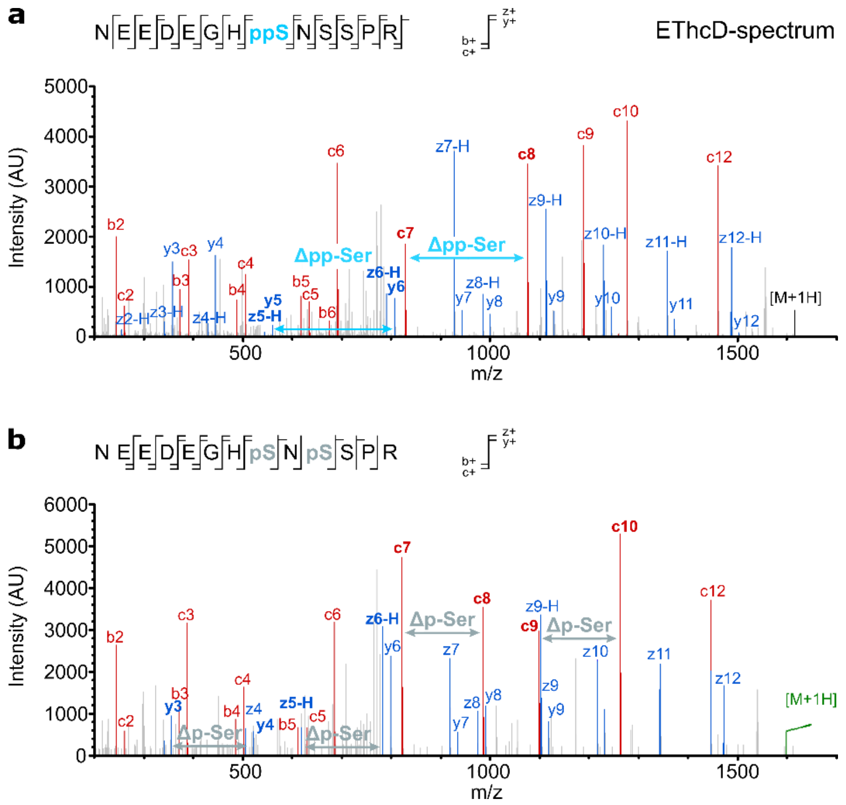


**Figure S1:** EThcD spectra of pyrophosphopeptide from figure 1c and corresponding bisphosphopeptide.

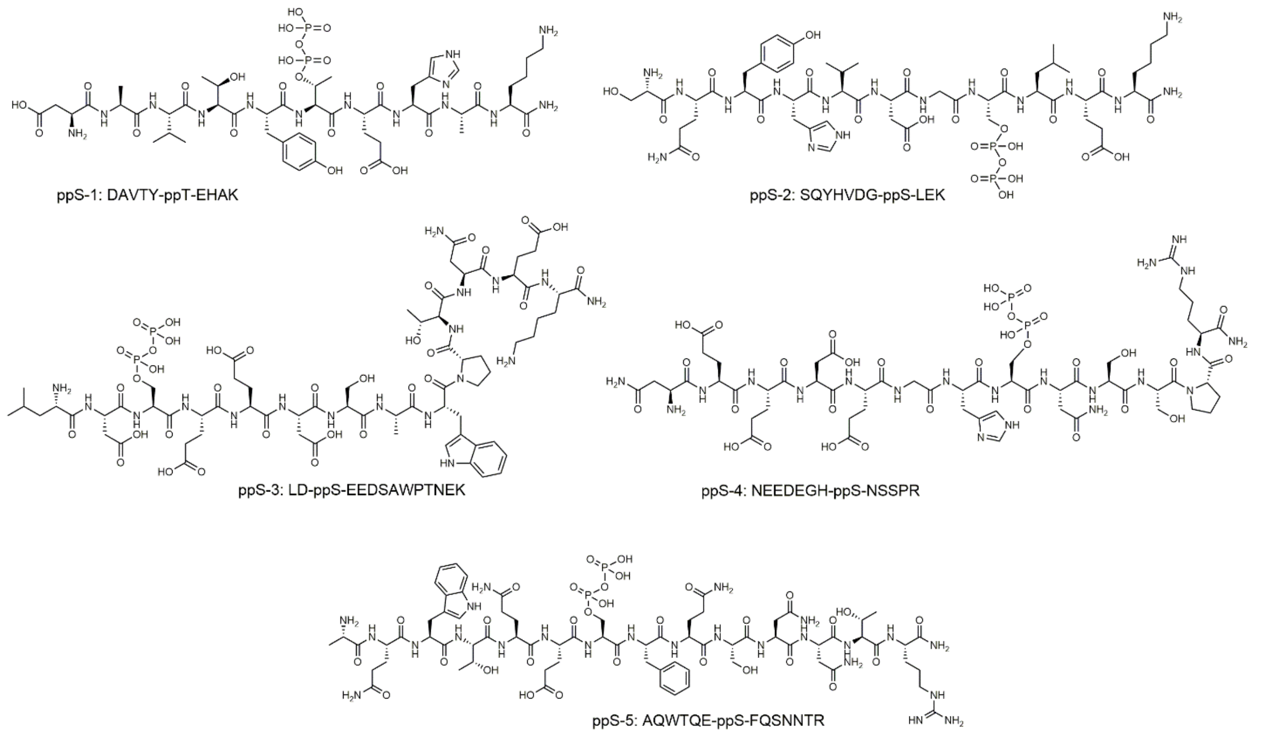


**Figure S 2:** Synthetic pyrophosphopeptides for validation of sample preparation workflow

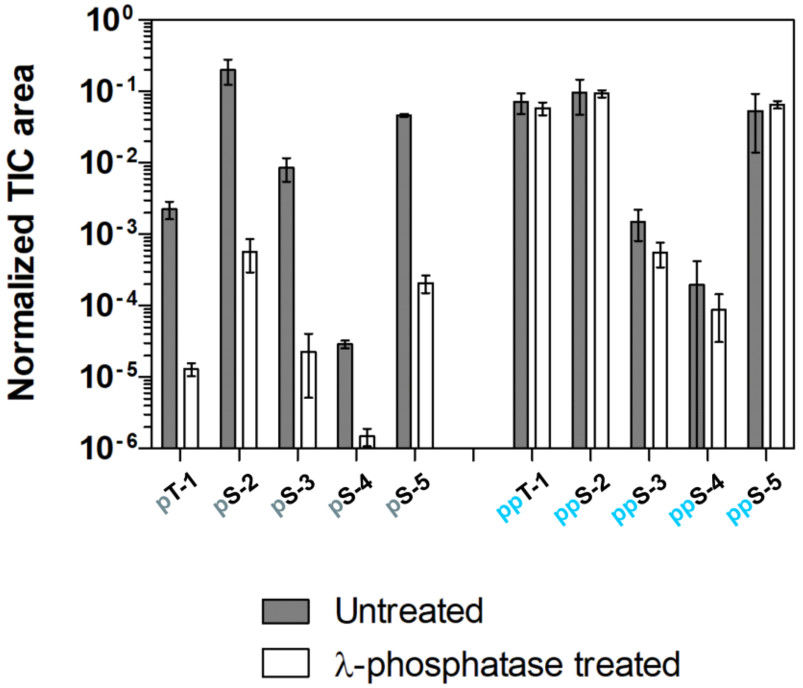


**Figure S3:** Pyrophosphopeptidepeptide stability to λ-phosphatase

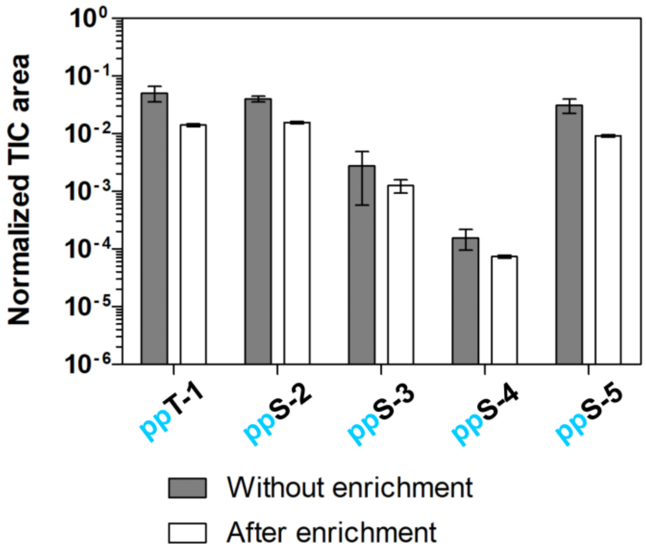


**Figure S4:** Showing retention of pyrophosphopeptides during SIMAC enrichment

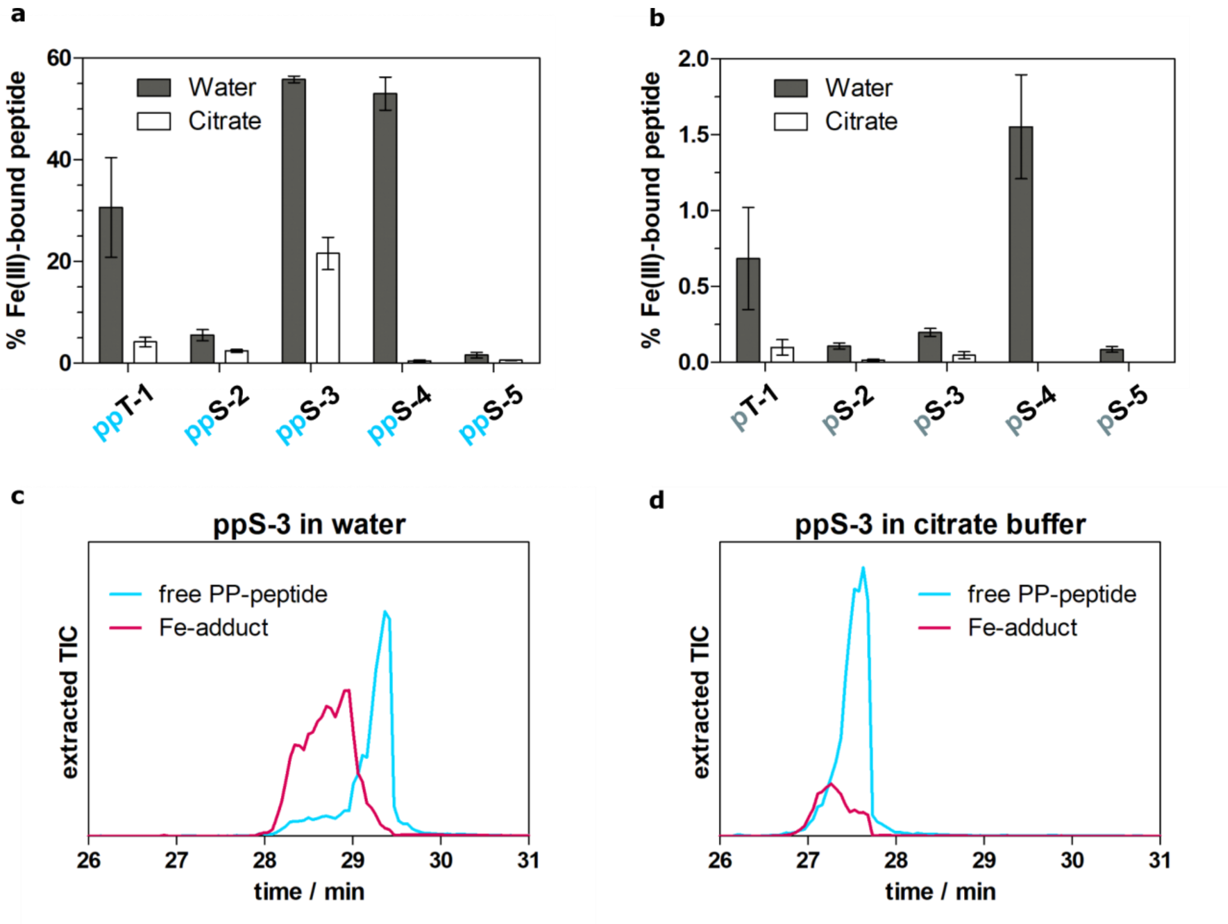


**Figure S5:** Improvement of LC-peak shape and reduction of Fe-adducts by citrate resuspension buffer

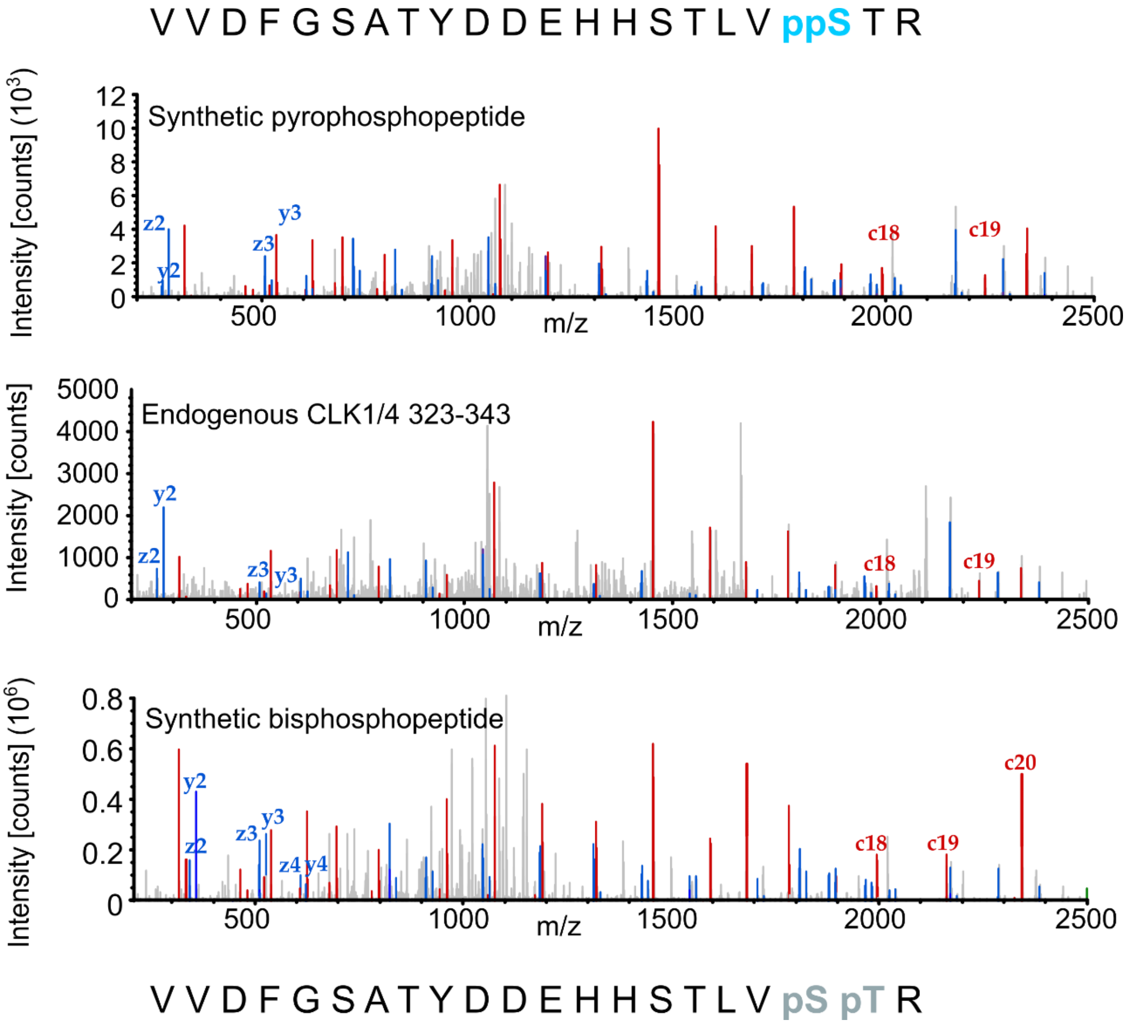


**Figure S6:** EThcD spectra of an endogenous pyrophosphopeptide (CLK1 323-343) and the corresponding synthetic pyrophospho- and bisphosphopeptide.

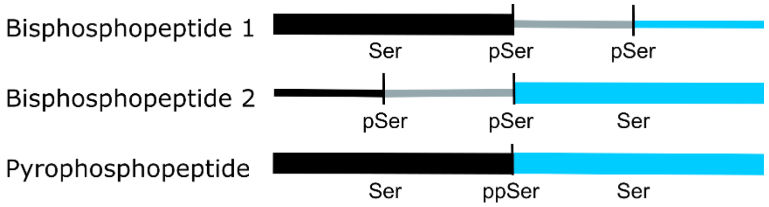


**Figure S7:** Schematic demonstrating how a mixture of bisphosphopeptides may contain all fragments typical of the corresponding pyrophosphopeptide and lead to false-positive assignment of a pyrophosphopeptide.

#### Nucleolar and coiled-body phosphoprotein 1 (NOLC1)

```
10         20         30         40         50
MADAGIRRVV PSDLYPLVLG FLRDNQLSEV ANKFAKATGA TQQDANASSL
        60         70         80         90        100
LDIYSFWLKS AKVPERKLQA NGPVAKKAKK KASSSDSEDS SEEEEEVQGP
       110        120        130        140        150
PAKKAAVPAK RVGLPPGKAA AKASESSSSE ESSDDDDEED QKKQPVQKGV
       160        170        180        190        200
KPQAKAAKAP PKKAKSSDSD SDSSSEDEPP KNQKPKITPV TVKAQTKAPP
       210        220        230        240        250
KPARAAPKIA NGKAASSSSS SSSSSSSDDS EEEKAAATPK KTVPKKQVVA
       260        270        280        290        300
KAPVKAATTP TRKSSSSEDS SSDEEEEQKK PMKNKPGPYS SVPPPSAPPP
       310        320        330        340        350
KKSLGTQPPK KAVEKQQPVE SSEDSSDESD SSSEEEKKPP TKAVVSKATT
       360        370        380        390        400
KPPPAKKAAE SSSDSSDSDS SEDDEAPSKP AGTTKNSSNK PAVTTKSPAV
       410        420        430        440        450
KPAAAPKQPV GGGQKLLTRK ADSSSSEEES SSSEEEKTKK MVATTKPKAT
       460        470        480        490        500
AKAALSLPAK QAPQGSRDSS SDSDSSSEE EEEKTSKSAV KKKPQKVAGG
       510        520        530        540        550
AAPSKPASAK KGKAESSNSS SSDDSSEEEE EKLKGKGSPR PQAPKANGTS
       560        570        580        590        600
ALTAQNGKAA KNSEEEEEEK KKAAVVVSKS GSLKKRKQNE AAKEAETPQA
       610        620        630        640        650
KKIKLQTPNT FPKRKKGEKR ASSPFRRVRE EEIEVDSRVA DNSFDAKRGA
       660        670        680        690
AGDWGERANQ VLKFTKGKSF RHEKTKKKRG SYRGGSISVQ VNSIKFDSE
```

#### Upstream binding factor 1 (UBF1)

```
10         20         30         40         50
MNGEADCPTD LEMAAPKGQD RWSQEDMLTL LECMKNNLPS NDSSKFKTTE
        60         70         80         90        100
SHMDWEKVAF KDFSGDMCKL KWVEISNEVR KFRTLTELIL DAQEHVKNPY
       110        120        130        140        150
KGKKLKKHPD FPKKPLTPYF RFFMEKRAKY AKLHPEMSNL DLTKILSKKY
       160        170        180        190        200
KELPEKKKMK YIQDFQREKQ EFERNLARFR EDHPDLIQNA KKSDIPEKPK
       210        220        230        240        250
TPQQLWYTHE KKVYLKVRPD ATTKEVKDSL GKQWSQLSDK KRLKWIHKAL
       260        270        280        290        300
EQRKEYEEIM RDYIQKHPEL NISEEGITKS TLTKAERQLK DKFDGRPTKP
       310        320        330        340        350
PPNSYSLYCA ELMANMKDVP STERMVLCSQ QWKLLSQKEK DAYHKKCDQK
       360        370        380        390        400
KKDYEVELLR FLESLPEEEQ QRVLGEEKML NINKKQATSP ASKKPAQEGG
       410        420        430        440        450
KGGSEKPKRP VSAMFIFSEE KRRQLQEERP ELSESELTRL LARMWNDLSE
       460        470        480        490        500
KKKAKYKARE AALKAQSERK PGGEREERGK LPESPKRAEE IWQQSVIGDY
       510        520        530        540        550
LARFKNDRVK ALKAMEMTWN NMEKKEKLMW IKKAAEDQKR YERELSEMRA
       560        570        580        590        600
PPAATNSSKK MKFQGEPKKP PMNGYQKFSQ ELLSNGELNH LPLKERMVEI
       610        620        630        640        650
GSRWQRISQS QKEHYKKLAE EQQKQYKVHL DLWVKSLSPQ DRAAYKEYIS
       660        670        680        690        700
NKRKSMTKLR GPNPKSSRTT LQSKSESEED DEEDEDDEDE DEEEEDDENG
       710        720        730        740        750
DSSEDGGDSS ESSSEDESED GDENEEDDED EDDDEDDDED EDNESEGSSS
       760
SSSSSGDSSD SDSN
```

**Figure S8: UBF1 sequence composition is distinctive from NOLC1.** The predicted IDRs on NOLC1 and UBF1 are indicated with a light blue highlight. The ppSer residues identified in NOLC1 are in turquoise, and the predicted ppSer residues in UBF1 are in blue. Lys or Arg residues marking the trypsin cleavage sites of the pyrophosphopeptides identified in NOLC1 are in red, and the trypsin cleavage sites neighboring the predicted pyrophosphosites in UBF1 are in magenta. The identified tryptic pyrophosphopeptides in NOLC1 are underlined, and the predicted pyrophosphopeptides in UBF1 are indicated with a dotted line.

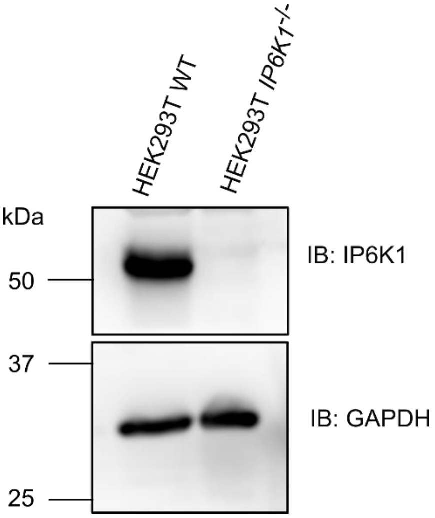


**Figure S9:** Western-Blot demonstrating the absence of IP6K1 in HEK293T knock-out cell line.

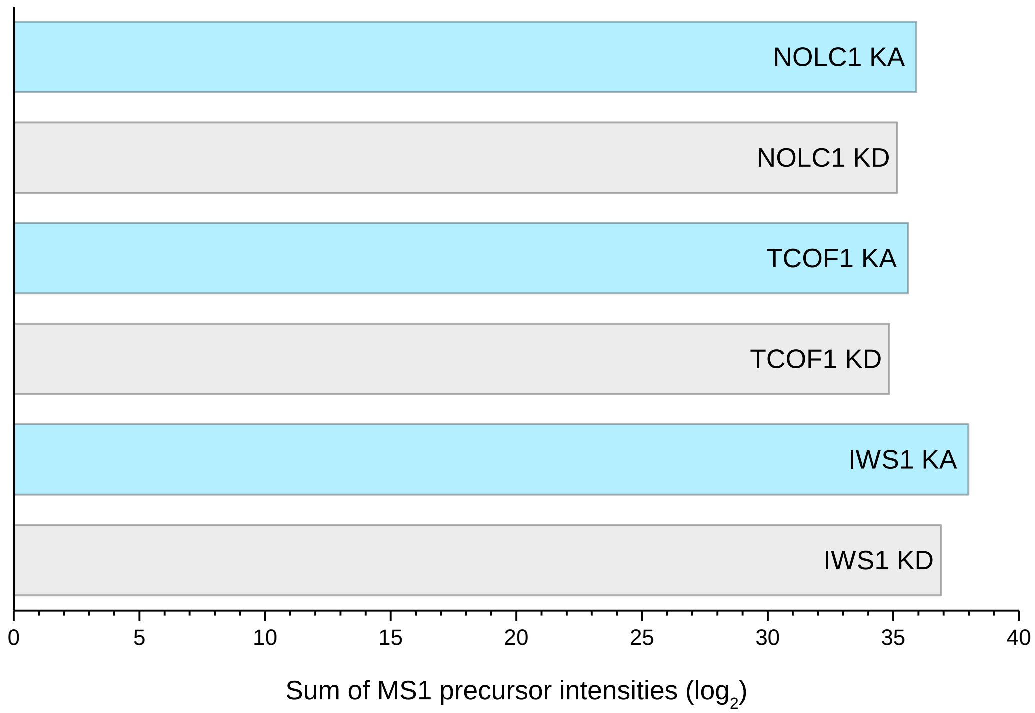


**Figure S10:** Sum of MS1 precursor intensities of all CID spectra recorded in pulldown-samples. Comparison between samples from a background with high (IP6K1-kinase active, KA) or low (IP6K1-kinase dead, KD) PP-InsP levels to demonstrate equal loading. For each of the three target proteins (NOLC1, TCOF1, IWS1), the sum of precursors is in the same order of magnitude for KA and KD conditions.

#### List of SI tables:

Table S1:

Sheet1: All sites from HEK293T triplicate and all sites from HCT116 method development, including IuPRED and Scansite analysis.

Sheet 2: All sites from HEK293T

Sheet 3: All sites from HCT116

Sheet 4: IuPred analysis of 71 proteins

Table S2: Comparison of sites from HEK293T triplicate run, including MS details.

Table S3: Complete GO results

Table S4: Pyrophosphosite prediction on FC-annotated proteins using IUPred2A and Scansite 4.0

Table S5: Details of pyrophosphosites found in pulldown samples

### Materials and methods:

#### Pyrophosphoproteomics sample preparation workflow

*Cell Lysis and digestion*

HEK293T cells (ATCC, CRL-3216) were cultured by seeding 2 x $10^6$ cells into eight 15 cm culture plates in DMEM (5% FBS, 1 mM L-Glu, 50 u/ml penicillin, 100 µg/ml streptomycin). Plates were cultured for three days, with medium exchange on day 3 and harvest on day 4 (80-90% confluency). To lyse, plates were washed twice with 5 ml 0.9% NaCl, then 2 ml lysis buffer (8 M urea, 75 mM NaCl, 50 mM Tris (pH 8.2), 1 mM NaF, 1 mM β–glycerophosphate, 1 mM sodium orthovanadate, 10 mM sodium pyrophosphate, 1 mM PMSF; 1 c0mplete® EDTA-free protease inhibitor tablet (Roche) per 10 ml)1 was added to seven plates. Plates were incubated at 4°C for 10-15 min. The remaining plate was trypsinized and cells counted using a BioRad cell counter (avg. cell number in HEK293T triplicate: 6.6 x $10^6$ cells per ml).
Lysed plates were scraped using a spatula and the combined lysate transferred into a 50 ml falcon tube. The lysate was subjected to sonication at 4°C (on ice, 50% output, 0.5 cycle rate, 5 x 30 s, 30 s rest between pulses). The lysate was centrifuged (3200 g, 10 min, 4°C), aggregated insoluble material on the surface was removed and the supernatant decanted from the cell pellet and retained. Supernatant protein concentration was determined by BCA assay (commercial kit by Thermofisher Scientific, avg. 112 mg).
To reduce and alkylate lysate proteins, DTT in MilliQ (5 mM concentration in sample) was added and the sample incubated at 37°C for 1 h. After incubation, the sample was cooled to RT. Iodoacetamide in MilliQ (14 mM concentration in sample) was added and the sample incubated at RT in the dark for 30 min. Remaining iodoacetamide was quenched with second aliquot of DTT (10 mM final concentration in sample), and the sample incubated for 15 min at RT in the dark.
To digest, sequencing grade modified trypsin (Promega, cat. V5111) was used. On ice, the lysate was diluted approx. 1:5 with 25 mM Tris (pH 8.0). CaCl2 (1 mM concentration in sample) was added. Trypsin was added at a ratio of approx. 1:50 protease to protein. The sample was incubated for 16 h at 37°C with 500 RPM agitation.

*Lysate desalting*

Neat TFA (approx. 0.4% of sample volume) was added (approx. pH 2), and the sample was centrifuged for 10 min at 3,200 g. The supernatant of the centrifuged protein solution was desalted using four SepPak® tC18 3 cc 500 mg cartridges (Waters, cat. WAT043425, loading of approx. 20 mg/ cartridge). Air pressure was used to speed up washing steps but was not used during the loading or elution steps. To prepare, columns were washed with MeCN (9 mL) and then with 3 mL of 50% MeCN in H2O with 0.5% AcOH. The columns were equilibrated with 0.1% TFA in H2O (9 mL) and then each loaded with the digested peptide samples in 0.4% TFA in H2O. The columns were washed with 0.1% TFA in H2O (9 mL) to desalt the peptides. The counter-ion was exchanged by a final wash with 0.5% AcOH in H2O (1 mL). Peptides were eluted with 6 mL of 50% MeCN and 0.5% AcOH in H2O. The eluent was lyophilized overnight to isolate tryptic digest as a white solid (avg. yield: 84 mg) *λ –Phosphatase treatment*

Lyophilized tryptic digest (50 mg) was dissolved in 10 ml of phosphatase reaction buffer (50 mM HEPES, 100 mM NaCl, 1 mM MnCl2, 2 mM DTT, 0.01% Brij 35). The solution was then treated with 50000 units λ –phosphatase and incubated at 37°C with 300 RPM agitation for 5 h. Neat TFA was added (approx.0.4% of sample volume, pH 2), and the sample centrifuged (10 min 3,200 g) to remove any cellular debris. The samples were then desalted as described in the previous step, except only three SepPak cartridges were used. The eluent was again lyophilized to isolate the tryptic digest as a white solid (avg. yield: 45 mg)

*SIMAC enrichment*

A High-select Fe-NTA Phosphopeptide enrichment kit from Thermofisher Scientific was used to enrich the pyrophosphopeptides using a protocol adapted from the manufacturer's instructions to facilitate sequential immobilized metal affinity chromatography (SIMAC)2.
Tryptic digest (40 mg) treated with λ-phosphatase from the previous step was dissolved in 800 µl of SIMAC loading buffer (0.1% TFA, 50% MeCN). Four columns were washed twice with 200 µl SIMAC loading buffer, then closed with a plug supplied with the kit and loaded with 200 µl of the digest solution (10 mg per column). The columns were incubated for 30 min at room temperature. Every ten minutes, the columns were gently tapped for 10 s to resuspend the Fe-NTA resin in the peptide solution.
After this binding step, columns were washed three times with 200 µl SIMAC loading buffer, once with 200 µl MilliQ water, twice with 100 µl SIMAC buffer A (1% TFA, 20% ACN, to selectively elute monophosphopeptides), and once again with 200 µl MilliQ water.
Elution was performed twice with 100 µl SIMAC buffer B (0.5% NH4OH in water), into clean 2 ml low protein bind Eppendorf tubes. The eluent from four columns was then combined into a single 2 ml protein low bind Eppendorf tube. This yielded an 800 µl peptide solution that was directly frozen using liquid nitrogen and lyophilized overnight.

*Fractionation*

After enrichment, the material was dissolved in 100 µl HPLC buffer A, spun at 21000 g for 10min to pellet any insoluble components and transferred to an HPLC vial for fractionation. Separation was achieved using a decreasing pH gradient to exploit pKa differences between acidic amino acid side chains, phosphoryl groups and pyrophosphoryl groups, with lower pKa peptides expected to retain longer. Additionally, a decreasing acetonitrile gradient was employed with the aim to rapidly elute peptides without significant negative charge in the early fractions of the separation. Fractionation was performed on an Agilent 1260 Infinity HPLC, equipped with an IonPac® AS24 2 mm analytical ultra-hydrophilic SAX column (Thermofisher Scientific) and the corresponding IonPac® AG24 guard column. HPLC buffers were prepared freshly.
HPLC buffer A: 20% MeCN in distilled water, 1% formic acid, pH 9.0 set with NH4OH; buffer B: 1% MeCN, 1% NH4OH, pH 2.8 set with formic acid. Flow rate: 0.2 ml/min. Gradient: 0-5 min, 100% A; 5-50 min, gradient increase to 100% B, 50-60 min, 100% B. Sample injection volume was 100 µl. Fractions were collected every 5 min, 12 1 ml fractions in total.

Fractions were transferred to 2ml protein low bind Eppendorf tubes and lyophilized overnight. The resulting solids were redissolved in 200 µl 20% MeCN and transferred to glass LC-MS vials, then dried using a vacuum centrifuge for 2h at RT. Samples were stored at -20°C until LC-MS analysis.

**Liquid chromatography combined with mass spectrometry (LC-MS):**

Fractions were re-suspended with 50 mM citric acid in 3 % acetonitrile solution before submitting to the LC-MS analysis. Sample separation was achieved by reverse phase (RP)-HPLC on a Thermo Scientific™ Dionex™ UltiMate™ 3000 system coupled on-line to an Orbitrap Fusion mass spectrometer (Thermo Fisher Scientific). For sample loading a PepMap C-18 trap-column (Thermo Fischer Scientific) of 0.075 mm ID x 50 mm length, 3 µm particle size and 100 Å pore size was used. Reversed-phase separation was performed using a 50 cm analytical column (in-house packed with Poroshell 120 EC-C18, 2.7 µm, Agilent Technologies) with mobile phase A contained 0.1% formic acid in water, and mobile phase B 0.1% formic acid in acetonitrile. The gradient started with 4% mobile phase B reaching 80% mobile phase B in 101 min, with total run time of 120 min including column wash and equilibration.

MS1 scans were acquired in the Orbitrap with a mass resolution of 120,000. MS1 scan range was set to 380 – 1400 m/z, with standard AGC target of 4e5, and maximum injection time of 50 ms. Precursor ions with charge states 2 - 4 were isolated with an isolation window of 1.6 m/z and dynamic exclusion of 15 sec. Precursor ions were selected with precursor priority to the higher charge state. MS2 scans were acquired in the Orbitrap with AGC target of 1e4 and maximum injection time of 100 ms. Precursor ions were fragmented using collision induced dissociation (CID) with a normalized collision energy of 25%. If neutral losses of 177.9432 Da from precursor ions were measured above a threshold of 15% of relative intensity in the CID scan, an additional spectrum of the same precursor ions was acquired using EThcD. EThcD spectra were measured in the Orbitrap with a resolution of 15,000 with AGC target of 1× 10e5 and maximum injection time of 2 s and normalized collision energy of 30%. Cycle time was set to 3 sec.

**Data analysis**

Raw files were analyzed using Proteome Discoverer (Thermofisher Scientific) version 2.4.
EThcD spectra were selected by spectrum selector, accepting precursor masses of 350 – 5000 Da.
Non-fragment filter was used as follows: Precursor ions were removed in a 1 Da window offset, and charge-reduced precursors and neutral losses within a 0.5 Da window offset.
The SequestHT node was used for peptide identification, searching against a full human proteome, digested by trypsin, with 2 missed cleavages allowed. Precursor mass tolerance was 10 ppm and fragment mass tolerance was 0.02 Da. Carbamidomethylation (C) was set as a static modification, and phosphorylation (S,T,Y), pyrophosphorylation (S,T) and Oxidation (M) were searched as dynamic modifications.
Peptide spectrum matches (PSMs) were filtered using the Fixed value PSM Validator, with a maximum accepted Delta Cn of 0.5.
*ptm*RS was used for automated PTM assignment. PhosphoRS mode was set to false, in order to detect phosphorylation and pyrophosphorylation in parallel.

**Validation of individual workflow steps:**

A set of five synthetic pyrophosphopeptides (***Figure S 2***) with a variety of sequence characteristics were produced as previously published.[3] Briefly, a phosphorimidazolide reagent was used to install a benzyl-protected β-phosphate onto a phosphoserine or phosphothreonine residue in a peptide, followed by Pd-catalysed hydrogenolysis of the protecting group and purification by C8-reverse phase HPLC. These standard peptides were used to validate individual steps of the sample preparation workflow.

*λ-phosphatase treatment:*

Five synthetic pyrophosphopeptides (20pmol each, see figure S2) and 0.5 µg HCT116 tryptic digest, were spiked into λ-phosphatase buffer (50 mM HEPES, 100 mM NaCl, 1 mM MnCl2, 2 mM DTT, 0.01% Brij 35). Total peptide amount was ca. 0.7 µg. λ-phosphatase was added (1 u/µg as in the proteomics workflow) and samples were incubated at 37°C, 300 RPM for 5 h in a heater shaker.
Four replicates containing PP-peptides, four negative controls without enzyme, four replicates containing the corresponding monophosphopeptides and four controls with monophosphopeptides but without enzyme were treated in this way.
All samples were desalted using C18-stage tips and subjected to LCMS-MS analysis as described above.
Extracted TICs of +2 precursor masses of the synthetic phospho- and pyrophosphopeptides were integrated on MS1 level and normalized against total background to achieve a relative quantification of peptide abundance in samples vs. controls.

*SIMAC enrichment:*

Four replicates containing 5 mg of a λ-phosphatase-treated tryptic digest from HCT116 cells and five synthetic pyrophosphopeptides (20 pmol each, see figure S2) and four controls containing only the HCT116 digest were dissolved in 200 µl SIMAC loading buffer per sample (0.1 % TFA, 50 % MeCN) and subjected to SIMAC enrichment as in the pyrophosphoproteomics workflow described above. After enrichment, 20 pmol of each synthetic peptide were added to each control sample (positive control). Eluates were lyophilized, cleaned using C18-stage-tips and dried by vacuum centrifugation. Peptides were redissolved in 6 µl injection buffer (50 mM sodium citrate, 3 % MeCN) and subjected to LC-MS analysis as described above.

Extracted TICs of +2 precursor masses of the synthetic phospho- and pyrophosphopeptides were integrated on MS1 level and normalized against total background to achieve a relative quantification of peptide abundance in samples vs. controls.

*Use of citrate resuspension buffer:*

Five synthetic pyrophosphopeptides (2 µl peptide mix, 20pmol each, figure S2) and 0.5µg HCT116 tryptic digest (1 µl) were mixed with either 3 µl water or 3 µl citrate buffer (100 mM sodium citrate, 6 % MeCN) and subjected to LC-MS analysis as described above.

Ion chromatograms of the most abundant free and iron-bound species $(M+Fe(III)-H)^{2+}$ and $(M+2H)^{2+}$ were extracted, integrated and normalized against total background to achieve a relative quantification of free versus iron-bound pyrophosphopeptides. The same experiment was performed with the corresponding monophosphopeptides.

**In-gel digestion after pulldown:**

Coomassie-stained bands were cut out, and washed with 200µl wash buffer (50 mM TEAB in 1:1 water:MeCN) until destained. In case of strongly colored bands, this process was repeated to fully remove Coomassie. The wash buffer was discarded and gel slices equilibrated for 10 min at 30°C in 50mM TEAB, the supernatant was discarded. 200µl MeCN were added to shrink and dry gel pieces. This step was repeated once more.
IWS1 samples were then reduced and alkylated. NOLC1 and TCOF1, having no Cys residues, were not. 100 µl 5 mM DTT were added to each tube and incubated for 45 min at 56°C, 300 rpm on a shaker; the supernatant was discarded. 100 µl 40 mM CAA in 50 mM TEAB were added and incubated for 30 min in the dark at room temperature; supernatant was discarded. Gel slices were incubated in 200 µl wash buffer for 5 min at 30°C, 300 rpm, then in 200 µl MeCN for 5 min. The MeCN step was repeated once more.

For digestion, 0.2 µg trypsin in 30µl 50 mM TEAB were added to each gel slice. In some cases, more buffer was added until the gel piece was covered. Samples were incubated overnight at 37°C, 300 rpm.
The next day, samples were briefly centrifuged, digestion was stopped by adding 30µl 0.5% TFA in MeCN. The supernatant was transferred to an MS vial. 20µl MeCN were added to the gel piece to shrink and dry it, then added into the same vial. The combined solutions were dried in a vacuum centrifuge and stored at -20°C until LC-MS analysis.
Samples were then redissolved in 10µl citrate injection buffer (50 mM sodium citrate, 3% MeCN) and sonicated for 5 min. 2 µl were injected and analysed in the same manner as the pyrophosphoproteomics samples described above.

**Expression constructs**

The following plasmids were used for expression of cDNA constructs in mammalian cells: N-terminally SFB (S-protein/FLAG/SBP)-tagged human TCOF1 (GenBank ID NM_000356.4) was a gift from Maddika Subba Reddy (Centre for DNA Fingerprinting and Diagnostics, Hyderabad); human UBF1 cDNA (GenBank ID NM_014233.4), a gift from Solomon Snyder (Johns Hopkins School of Medicine, Baltimore, USA) was subcloned into pCMV-Myc-N plasmid for expression with an N-terminal myc tag; full-length human IWS1 (GenBank ID NM_017969.3) was amplified using cDNA from HepG2 cells and cloned into N-terminal GFP or N-terminal SFB-tagged destination vectors using the Gateway cloning strategy (Thermo Fisher Scientific); N-terminally myc-tagged human NOLC1 cDNA (GenBank ID NM_001284388.2) expression plasmid was obtained from Sino Biological (HG16317-NM). The generation of catalytically active and inactive versions of C-terminally V5-tagged human IP6K1 (GenBank ID NM_001242829.2) has been described earlier.[4]

**Cell lines and transfection**

All cell lines used in this study were grown in Dulbecco's Modified Eagle Medium (DMEM) supplemented with 10% fetal bovine serum, 1 mM L-glutamine, 100 U/mL penicillin, and 100 µg/mL streptomycin in a humidified incubator with 5% CO2. Plasmid Midi kit (Qiagen) was used to purify all plasmids used for transfection, and cells were transfected using polyethylenimine (PEI) (Polysciences) at a ratio of 1:3 (DNA:PEI). Cells were harvested 48 h post-transfection for further analyses. Wild type and IP6K1-/- IP6K2-/- double knockout HCT116 cells were a gift from Adolfo Saiardi, University College London, London.[5] The HEK293T cell line was used to generate IP6K1 knockout cells using the CRISPR-Cas9 strategy. A sgRNA sequence targeting exon 5 of IP6K1 (5′-CCTTGGACTCGGAGCGCATG-3′) was designed using the online tool Benchling (https://www.benchling.com), and cloned into pU6-2A-GFP-2A-Puro plasmid (a gift from P. Chandra Shekar, Centre for Cellular & Molecular Biology, Hyderabad) which co-expresses Cas9. The plasmid was transfected into HEK293T cells using PEI, and after 48 h the transfected cells were selected using 2 µg/mL puromycin for 3 days. Serial dilution was performed to isolate single cell-derived colonies, which were screened for frameshift mutations by genotyping using a 3500xL Genetic Analyzer (Applied Biosystems). Primers used for genotyping were: FAM-tagged forward primer, 5′-GCAAAGAGTCCGAAGGTGGA-3′ and reverse primer, 5′-GAGGGGCAACGAGGATACTG-3′. Positive clones were sequenced to confirm mutations in exon 5, and the knockout was confirmed by immunoblot analysis with an IP6K1-specific antibody.

**Immunofluorescence**

Cells seeded on 12 mm glass coverslips in a 24-well plate were fixed with 4% paraformaldehyde for 15 min and permeabilized using PBS containing 0.15% Triton X-100 (PBST) for 15 min at room temperature. The cells were incubated with blocking solution (5% BSA in PBST) for 1 h at room temperature to block non-specific antibody binding, followed by incubation with primary antibodies diluted in the blocking solution overnight at 4°C. Cells were washed thrice with PBST and incubated with fluorophore-conjugated secondary antibodies (Alexa Fluor 488 or Alexa Fluor 568 goat anti-rabbit or anti-mouse IgG; Thermo Fisher Scientific, 1:500) for 1 h in the dark at room temperature. Post incubation, the cells were washed thrice with PBST and mounted on glass slides, using an antifade mounting medium with DAPI (Vecta Labs). Images in Fig 4b were captured on a Leica TCS SP8 confocal microscope equipped with 405, 488, 514, 561, and 633 nm lasers using a 63x 1.4 NA oil immersion objective. For colocalization studies in Figs. 4c, d, images were acquired using the Elyra 7 structured illumination microscopy (SIM) module of the Zeiss LSM 980 confocal microscope, equipped with 405, 488, 561, and 642 nm lasers, and 63x 1.4 NA oil immersion objective. All immunofluorescence images are in z-stacks and are shown as maximum intensity projections using LAS X (Fig. 4b) or ZEN (Figs. 4c, d) software.

**Protein pull-down and immunoblotting**

Cells were harvested 48 h post-transfection, and lysed for 1 h at 4°C on an end-over-end mixer in 1% lysis buffer (50 mM HEPES, pH 7.4, 150 mM NaCl, 1% Nonidet P-40, 1 mM EDTA, protease and phosphatase inhibitor cocktail) followed by sonication for 10 sec at 30% amplitude (SONICS, VCX750). To the lysate, the specific antibody was added and incubated overnight at 4°C on an end-over-end mixer. The antibody bound protein was pulled down using Protein A or Protein G Sepharose beads (GE Healthcare) for 1 h. The beads were washed three times with lysis buffer and used for [$\beta^{32}$P]5PP-InsP$_5$-mediated pyrophosphorylation. For SFB-tagged proteins, streptavidin sepharose beads (GE Healthcare) were added to the cell lysate and incubated overnight at 4˚C prior to washing and pyrophosphorylation with [$\beta^{32}$P]5PP-InsP$_5$. For MS analysis of overexpressed proteins (Fig. 5a) 4% of cell lysate was used for immunoblotting and 90% was pulled down on streptavidin sepharose beads. The beads were washed with lysis buffer, boiled in 1× Laemmli buffer, and proteins were resolved by SDS-PAGE. The bands were visualized with 0.2% Coomassie brilliant blue R-250, excised, digested with trypsin and subjected to neutral loss-triggered EThcD MS as described above. For immunoblotting, proteins were transferred to a PVDF membrane, and detected using standard western blotting techniques with protein- or tag-specific antibodies. Chemiluminescence was detected using the GE ImageQuant LAS 500 imager.

**Protein pyrophosphorylation using [β$^{32}$P]5PP-InsP$_5$**

Radiolabeled [β$^{32}$P]5PP-InsP$_5$ synthesis was conducted as described earlier[6], with a few modifications. Briefly, for a 200 µL reaction, 200 µM inositol hexakisphosphate (InsP$_6$, SiChem) was incubated for 6 h at 37ºC with 3 mCi [γ32P]ATP and 50 µM unlabeled ATP in the presence of 80 ng/µL of purified Entamoeba histolytica IP6 kinase (IP6KA) in buffer containing 100 mM MES pH 6.8, 30 mM MgSO4, 250 mM NaCl and 5 mM DTT. [β32P]5PP-InsP$_5$ was purified by strong anion exchange HPLC (Waters) as described earlier.[6]
Proteins were subjected to phosphorylation by CK2, and pyrophosphorylation by [β32P]5PP-InsP$_5$ as described earlier[7–9]. For in vitro pyrophosphorylation assays, immunoprecipitated proteins on beads were first pre-phosphorylated with CK2 (New England Biolabs) in protein kinase buffer (New England Biolabs) and 0.5 mM Mg2+-ATP for 30 min at 30°C. Beads were washed in cold PBS and incubated in pyrophosphorylation buffer (25 mM HEPES, pH 7.4, 50 mM NaCl, 6 mM MgCl2, and 1 mM DTT) containing 3-5 µCi [β32P]5PP-InsP$_5$ at 37°C for 15 min. LDS sample buffer (Thermo Fisher Scientific) was added to the beads, the sample was heated at 95°C for 5 min, resolved on a 4–12% NuPAGE Bis-Tris gel (Thermo Fisher Scientific), and transferred to a PVDF membrane (GE Life Sciences). Pyrophosphorylation was detected using a phosphorimager (Typhoon FLA-9500), and proteins were detected by immunoblotting. To improve visualisation, the phosphorimager scan and immunoblots were subjected to uniform "Levels" adjustment in Adobe Photoshop.
Back-pyrophosphorylation was conducted as described earlier[9–11] and the method is explained schematically in Fig 5b. Overexpressed or endogenous UBF1 was immunoprecipitated from IP6K1-/- HEK293T cells expressing either active or kinase-dead IP6K1. The immunoprecipitated proteins were subjected to pyrophosphorylation as described above. Radiolabeled protein as a fraction of total immunoprecipitated protein was quantified using Fiji software[12].

**RT-qPCR**

Cells were lysed using TRIzol reagent (Thermo Fisher Scientific), and total RNA was extracted using a kit (HiMedia). Where indicated, cells were treated with 10 µM TNP [13], 1 µM SC-919 [14], or DMSO as a vehicle control for 5 h before RNA isolation. 2 µg RNA was used to prepare cDNA by reverse transcription with SuperScript Reverse Transcriptase III (Thermo Fisher Scientific) using random hexamers. Two sets of 45S pre-rRNA specific primers were designed to amplify the region between sites 01 (also called A') and A0 on the human 47S pre-rRNA transcript[15]. GAPDH transcript was used as an internal control. The sequences of the primers used were: Primer Set 1 forward 5′-CCTTCCCCAGGCGTCCCTCG-3′ and reverse 5′-GGCAGCGCTACCATAACGGA-3′; Primer Set 2, forward 5′-GAACGGTGGTGTGTCGTT-3′ and reverse 5′-GCGTCTCGT-CTCGTCTCACT-3′; GAPDH forward, 5′-ACATCAAGAAGGTGGTGAAGCA-3′ and reverse 5′-TTGTCATACCAGGAAATGAGCTTG-3′. qPCR was conducted with SYBR™ Green PCR Master Mix on CFX96 Touch Real-Time PCR Detection System (Biorad). All samples were run in technical duplicates. The fold change (ΔΔCT) method[16] was used to calculate the difference in transcript levels. ΔCT refers to the CT value for the target pre-rRNA normalized to the CT value for GAPDH in the same sample. ΔΔCT values were determined as a relative change in ΔCT in HCT116 DKO compared to WT cells or drug treated compared to control cells. 2-ΔΔCT was used to represent fold changes.

#### Statistical analysis

Statistical analyses and graph preparation were done using GraphPad Prism 8. The immunofluorescence analyses were conducted over three independent experiments and representative images are shown in Figs. 4b), c). The in vitro pyrophosphorylation assays were conducted as at least three independent experiments for each protein, and representative images are shown in Figs. 4f)-i). Back-pyrophosphorylation data was compiled from at least three independent experiments and representative images are shown in Figs. 5c), d). Neutral loss-triggered EThcD MS analysis was conducted on overexpressed proteins in one or two experimental sets. RT-qPCR data was compiled from three biological replicates. Quantified data are presented as mean ± SEM for the indicated number of biological replicates (N). P values were calculated using a one-sample t-test. $P \leq 0.05$ was considered statistically significant.

13. Padmanabhan, U., Dollins, D. E., Fridy, P. C., York, J. D. & Downes, C. P. Characterization of a selective inhibitor of inositol hexakisphosphate kinases. Use in defining biological roles and metabolic relationships of inositol pyrophosphates. *J. Biol.*

*Chem.* **284**, 10571–10582 (2009).

14. Moritoh, Y. *et al.* The enzymatic activity of inositol hexakisphosphate kinase controls circulating phosphate in mammals. *Nat. Commun.* **12**, (2021).

15. Mullineux, S.-T. & Lafontaine, D. L. J. Mapping the cleavage sites on mammalian pre-rRNAs: Where do we stand? *Biochimie* **94**, 1521–1532 (2012).

16. Schmittgen, T. D. & Livak, K. J. Analyzing real-time PCR data by the comparative CT method. *Nat. Protoc.* **3**, 1101–1108 (2008).
